# Mapping brain-wide excitatory projectome of primate prefrontal cortex at submicron resolution: relevance to diffusion tractography

**DOI:** 10.1101/2021.09.13.460040

**Authors:** Mingchao Yan, Wenwen Yu, Qian Lv, Qiming Lv, Tingting Bo, Xiaoyu Chen, Yilin Liu, Yafeng Zhan, Shengyao Yan, Xiangyu Shen, Baofeng Yang, Zilong Qiu, Yuanjing Feng, Xiaoyong Zhang, He Wang, Fuqiang Xu, Zheng Wang

**Affiliations:** Institute of Neuroscience, State Key Laboratory of Neuroscience, Center for Excellence in Brain Science and Intelligence Technology, Chinese Academy of Sciences, Shanghai 200031, China; University of Chinese Academy of Sciences, Beijing100049, China; Institute of Science and Technology for Brain-inspired Intelligence, Fudan University, Shanghai 200433, China; College of Information Engineering, Zhejiang University of Technology, Hangzhou, 310014, China; Shenzhen Key Lab of Neuropsychiatric Modulation and Collaborative Innovation Center for Brain Science, Guangdong Provincial Key Laboratory of Brain Connectome and Behavior, CAS Key Laboratory of Brain Connectome and Manipulation, Brain Cognition and Brain Disease Institute (BCBDI), Shenzhen Institutes of Advanced Technology, Shenzhen-Hong Kong Institute of Brain Science-Shenzhen Fundamental Research Institutions, Shenzhen 518055, China

**Author notes:** These authors contributed equally to this work (Mingchao Yan, Wenwen Yu). **Corresponding author:** Zheng Wang (Institute of Neuroscience, Chinese Academy of Sciences, 320 Yueyang Road, Shanghai 200031, China) **Email:**.

## Abstract

Resolving trajectories of axonal pathways in the primate prefrontal cortex remains crucial to gain insights into higher-order processes of cognition and emotion, which requires a comprehensive map of axonal projections linking demarcated subdivisions of prefrontal cortex and the rest of brain. Here we report a mesoscale excitatory projectome issued from the ventrolateral prefrontal cortex (vlPFC) to the entire macaque brain by using viral-based genetic axonal tracing in tandem with high-throughput serial two-photon tomography, which demonstrated prominent monosynaptic projections to other prefrontal areas, temporal, limbic and subcortical areas, relatively weak projections to parietal and insular cortices but no projections directly to the occipital lobe. In a common 3D space, we quantitatively validated an atlas of diffusion tractography-derived vlPFC connections with correlative enhanced green fluorescent protein-labelled axonal tracing, and observed generally good agreement except a major difference in the posterior projections of inferior fronto-occipital fasciculus. These findings raise an intriguing question as to how neural information passes along long-range association fiber bundles in macaque brains, and call for the caution of using diffusion tractography to map the wiring diagram of brain circuits.

## Introduction

Higher-order processes of cognition and emotion regulation that depend on the prefrontal cortex are all based on multiple, long-range connections between neurons^1, 2, 3^. Axons connecting local and distant neurons form a fundamental skeleton of the brain circuitry, which is of paramount importance to fathom the organization of in-/output pathways that enable those vital functions^4, 5^. Given the complexity and heterogeneity of the primate prefrontal cortex^2^, understanding the working mechanisms of the prefrontal cortex requires a comprehensive map of axonal projections linking its demarcated subdivisions and the rest of brain. A subdivision of the prefrontal cortex - the ventrolateral section (vlPFC), which mainly spans Brodmann’s Areas 44, 45a/b, 46v/f and 12r/l^6^, is central to a variety of functions including language, objective memory and decision making^7, 8^. Emerging evidence further demonstrates abnormalities of vlPFC in tight association with deficits in cognitive flexibility^1, 9, 10^, suggesting that an elaborate delineation of its hard wiring would shed light on the underlying neuropathology of psychiatric disorders^11^.

Such neuroanatomical inter-areal connectivity has been probed using invasive bulk injections of tracers and noninvasive imaging methods with millimeter-scale spatial resolution^12, 13, 14^. Histological neural tracing has been historically utilized for circuit/pathway mapping and continues to be the most reliable way of survey all myelinated axons in mammalian brains^12, 20, 21^, which has also been used as a gold standard to validate other modalities like diffusion tractography^18, 22, 23, 24, 25, 26^. Diffusion tractography, which has been developed in 1990s to estimate the tissue microstructure by means of spatial encoding of water molecule movements^15^, represents the only methodology capable of inferring the ensemble of anatomical connections in the living animal or human brain^16, 17^. But this technique is an indirect observation with limited resolution and accuracy, and its reliability of false negative and false positive findings remains to be fully validated in a 3D space ^18, 19^. Notably, some classic tract tracing methods are not sensitive to specific neuronal types or axonal trajectories. They do not report the traveling course in a 3D space through which the axons travel for a remarkably long distance (i.e., over centimeter length). The pursuit of long-range axonal fiber tracing across the entire monkey brain has become feasible thanks to rapid advance in viral and genetic tools in the primate species, tissue labeling, large-scale microscopy and computational image analysis^27, 28, 29, 30^. Moreover, viral-based techniques for targeting specific neuronal types in macaque brain have achieved remarkable success^27, 31^, which may furnish the requisite biological detail including excitatory and inhibitory in-/output to enrich structural network reconstructions for improved prediction of brain function^32^. However, it remains unclear thus far what type of viral vector is suitable for long-range axonal fiber tracing^12, 20^.

In the present study, we aim to establish a comprehensive brain-wide excitatory projectome of the vlPFC in macaque monkeys using viral-based genetic tracing in tandem with serial two-photon (STP) tomography, a technique that has successfully achieved high-throughput fluorescence imaging of the entire mouse brain by integrating two-photon microscopy and tissue sectioning ^33^. In addition, in a common 3D space reconstructed with STP tomography, we intended to make a direct comparison of this mesoscale projectome to that derived from ultra-high field diffusion tractography (Fig. 1). Note that cross-comparison of the fiber details generated by two modalities with spatial scale differences in order of magnitude is technically demanding as many cellular structures or fiber pathways of biological interest are rather small relative to the voxel size of most diffusion MRI data^17^. One of challenging undertakings is to image long-range axonal fibers of many neurons with sufficiently high resolution to enable tracking axonal trajectories across the entire brain^33, 34^, which has stirred some debates such as right-angle fiber crossings^16, 24^ and the existence of the inferior fronto-occipital fasciculus in the primate brain^35^.

**Fig. 1.**
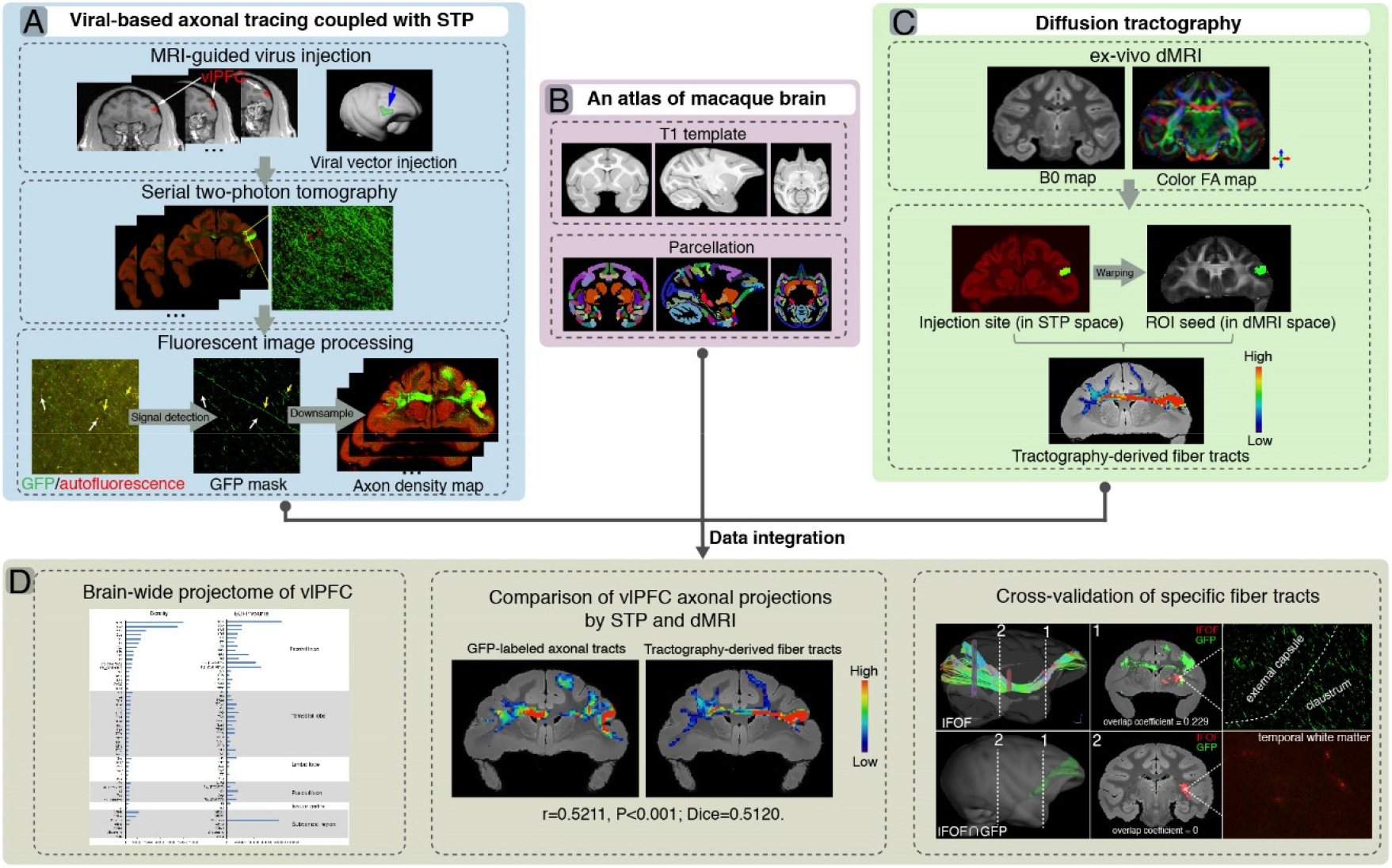
A flowchart diagram for brain-wide analyses of vlPFC projectome in macaques. The pipeline integrates the STP data in the mesoscopic domain (A) with macroscopic dMRI data (C) in a common 3D space (B). (**A**) T1 images were used to guide stereotaxic injection of AAV vectors to vlPFC (upper panel). High-throughput fluorescent images of viral-based genetic axonal tracing were acquired by STP tomography throughout the brain, which enables a close-up view and quantitative analysis of any region-of-interest (middle panel). A supervised machine learning approach was used for segmentation of GFP-positive signal and removal of autofluorescence in STP data. The serial segmented GFP images were down-sampled to compute the total signal intensity for each 200 μm × 200 μm grid by summing the number of signal-positive pixels in that grid and to generate the axonal density map (bottom panel). (**B**) An MRI atlas of cynomolgus macaques was used to construct a common 3D space. (**C**) Ex-vivo dMRI of macaque brain were acquired with using an 11.7T MRI scanner, illustrated as representative B0 (left) and direction-encoded color FA maps (right). Using the injection site identified from the STP data as seed regions, the target fiber tracts can be derived from diffusion tractography. (**D**) Integration of STP and dMRI data was implemented in a common 3D space, which allows quantitative analyses including whole-brain analysis of axonal projectome (left), comparison of vlPFC projectome by STP and dMRI (middle), and cross-validation of fiber tracking in both STP and dMRI (right).

## Results

### Determination of viral vectors for long-range anterograde tracing

We tested whether VSV, lentivirus, and AAV vectors with demonstrated success in rodents worked in the macaque brain and which vector was best suitable for long-range axonal fiber tracing. Five days after infection with VSV-ΔG, the neuronal cell bodies in the cerebral cortex (Fig. 2A and B) and thalamus (Fig. S1A) were clearly labelled with GFP, although only proximal neurites were labeled with no long-range axonal fibers detected (Fig. S2A). When the infection time was extended to about a month, we observed widespread axon loss and neuronal cell death (Fig. S1B-G). The infected neurons underwent morphological changes such as membrane blebbing (Fig. S1B and C), a key morphological change associated with the induction of apoptosis. Local injection with VSV-ΔG mediated rapid and transient gene expression nearby the injection site, and an extension of infection time evidently caused fatal neurotoxicity.

**Fig. 2.**
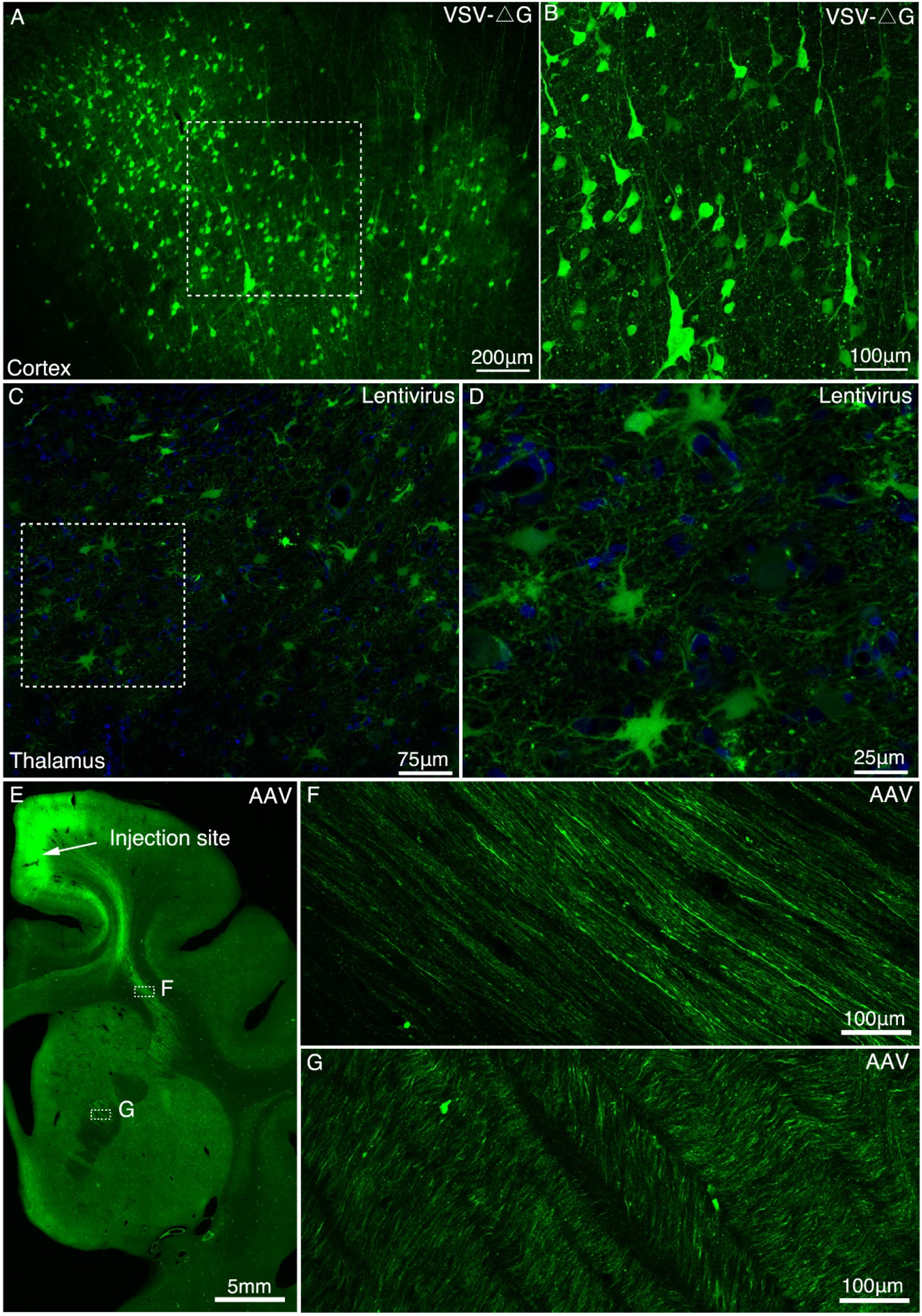
Determination of viral vectors for long-range anterograde tracing in macaques. (**A**) GFP-labeled neurons were found in the premotor cortex ∼5 days after injection of VSV-ΔG encoding Tau-GFP. (**B**) A magnified view illustrating the morphology of GFP-labeled neurons in the area outlined with a white box in (A). (**C**) Lentivirus construct was injected into the macaque thalamus and examined for transgene expression after ∼9 mouths. (**d**) High power views of the dotted rectangle in panel C. (**E**) GFP-labeled neurons and axons were observed in the premotor cortex ∼42 days after injection of AAV2/9 encoding Tau-GFP. Two dashed line boxes enclose the regions of interest: frontal white matter and ALIC, whose GFP signal are magnified in (**F**) and (**G**), respectively.

Lenti-Ubic-GFP exhibited stable expression in the cell soma even after 9 months (Fig. 2C and D), despite sparse labeling of GFP positive axons (Fig. S2B). By contrast, six weeks after AAV2/9-CaMKIIa-Tau-GFP was injected into the premotor cortex (Fig. 2E), axonal fiber bundles like anterior limb of internal capsule (ALIC) (Fig. 2G) were clearly visualized over several centimeters along the frontal white matter. As a validation test, AAV2/9 construct encoding mCherry was co-injected with AAV2/9 construct encoding Tau-GFP into the premotor cortex. And we found that the signal intensity of most Tau-GFP labeled axons was consistently higher than that of mCherry labeled axons (Fig. S3A-D).

We compared the axonal fiber tracing efficiency of VSV-ΔG, Lentivirus and AAV2/9 (AAV2/9-CaMKIIa-Tau-GFP) in the mediodorsal (MD) thalamus (Fig. S2). The density of axonal fibers labelled by AAV2/9 (Fig. S2C) was significantly higher (p < 0.001, Fig. S2D) than by Lentivirus (Fig. S2B), and VSV-ΔG (Fig. S2A). VSV-ΔG labeled axons sparse in the proximal, and the axonal density decreased sharply (Fig. S2D). Axons labeled by Lentivirus (Fig. S2B) were also significantly denser (p < 0.01, Fig. S2D) than by VSV-ΔG (Fig. S2A) distant from the injection site.

### Brain-wide excitatory projectome of vlPFC in macaques

AAV2/9 encoding Tau-GFP under the control of excitatory promoter CaMKIIa was determined as an anterograde tracer for mapping the excitatory projectome of vlPFC. The injection site in vlPFC, validated by STP images, including area 45, 12l and 44, was precisely located in cortical gray matter (Fig. 4A-D). To identify the cell type specificity of Tau-GFP gene expression driven by the CaMKII-α promoter, immunofluorescent staining was performed with antibodies against the excitatory neuron specific marker CaMKII-α and the inhibitory neuron-specific neurotransmitter GABA. GFP-positive neurons in the injection site were observed positive for CaMkIIa (Fig. S4A-C) and negative for GABA (Fig. S4D-F), indicating that the AAV labeled neurons were glutamate excitatory neurons.

**Figure 3.**
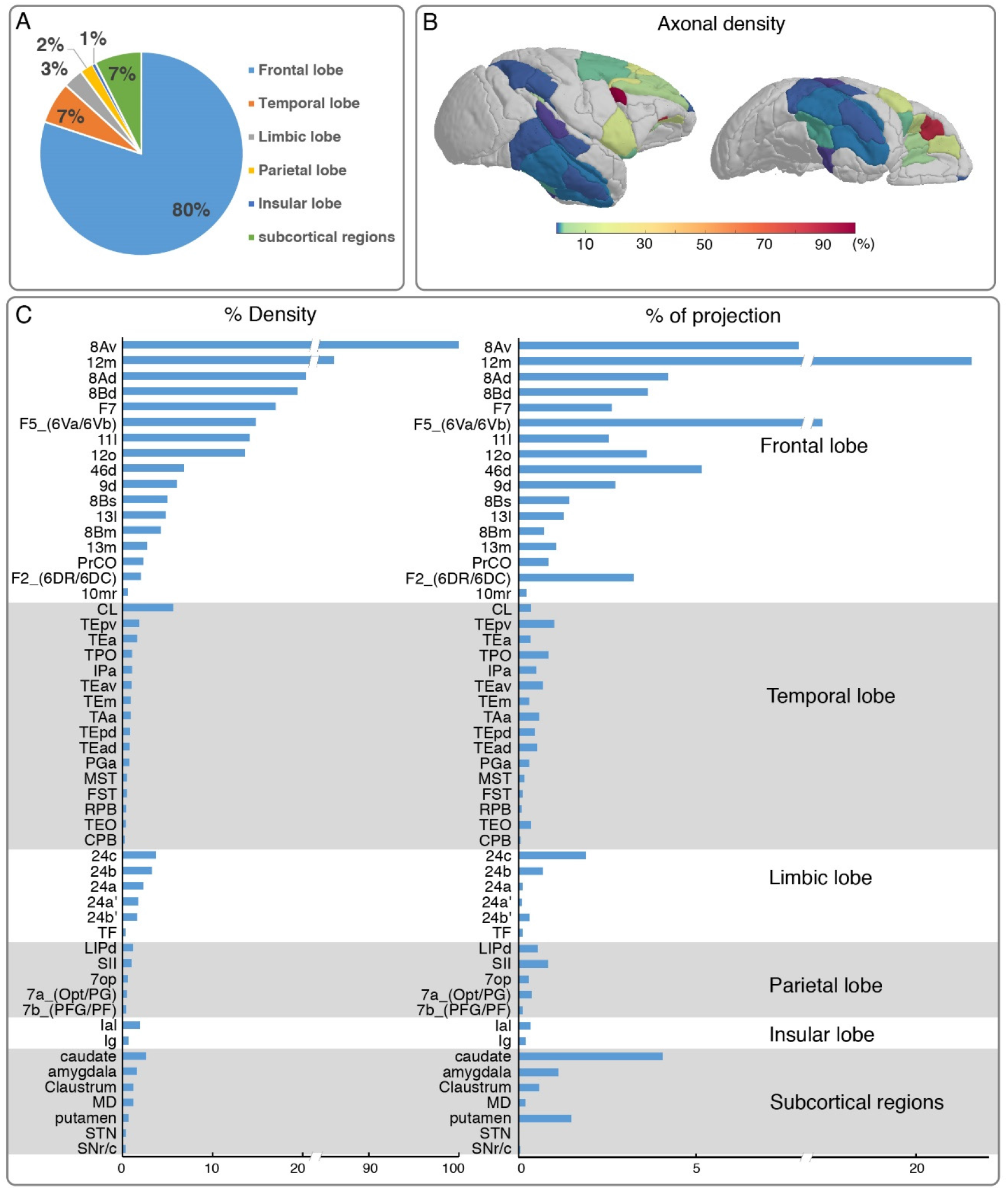
Brain-wide distribution of GFP-labelled excitatory projectome of vlPFC. (**A**) A pie chart shows the brain-wide distribution of vlPFC axonal projectome. (**B**) The normalized percentage distribution of axonal density was rendered onto a 3D brain surface. (**C**) The histogram plots show the vlPFC projections to other regions where the connectivity strength was quantified by the density of GFP-positive axons and proportion of total projection. We calculated the innervation density, given in percent of strongest projection.

**Figure 4.**
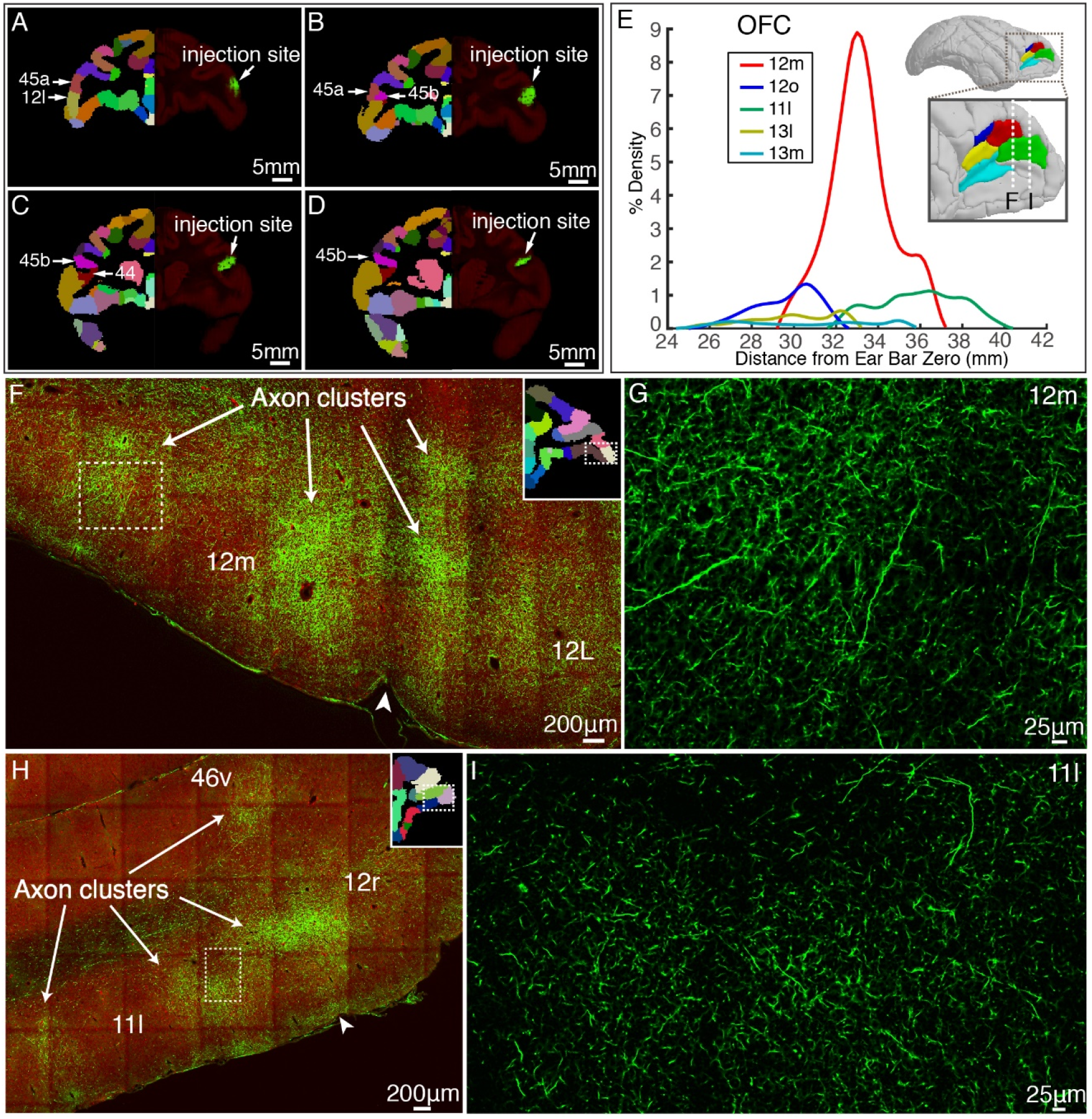
vlPFC projectome within the prefrontal lobe. (**A**-**D**) Representative coronal slices of the injection sites in vlPFC are shown overlaid with the monkey brain template (left hand side), mainly spanning areas 45a, 45b, 12l and 44. (**E**) Percentage of output density of vlPFC projectome along the anterior-posterior axis of the OFC. The inset shows the spatial location of individual Broadmann areas in OFC. Dotted lines indicate anterior-posterior position of the following fluorescent images. (**F**-**I**) Representative two-photon images of vlPFC axonal projections to OFC: 12m and 11l. Arrows indicate the axon clusters. Insets show the low power images of the section indicating the position of the higher power images.

To acquire a detailed account of the brain-wide vlPFC projectome, we analyzed its connectivity profile with other 173 parcellated regions in the monkey brain atlas using the STP tomography data (Fig. 3 and 4). The GFP-labelled projecting axons largely encompassed the anterior part of the brain including the frontal lobe, temporal lobe, limbic lobe, insular, and some subcortical regions, but no labeled axons were found in the occipital lobe (Fig. 3A-C). Within the frontal lobe, GFP-labeled projecting axons were markedly dense in the OFC, rostrally distributed in area 12m (Fig. 4E, F and G), 12o (Fig. 4E), 11l (Fig. 4E, H and I), 13l (Fig. 4E) and 13m (Fig. 4E). The 12m received strongest innervation from vlPFC relative to other OFC subregions (Fig. 4E). Laterally, axonal projections were found in the FEF including 8Av (Fig. S5A, B and C) and 8Ad (Fig. S5A and D). Dorsally, there were dense axonal fibers in the dorsal prefrontal cortex, including area 8Bd (Fig. S5E, F and G), 8Bm (Fig. S5E), 8Bs (Fig. S5E), 46d (Fig. S5E, J and K), and 9d (Fig. S5E, H and I). The 8Bd and 46d received relatively more innervation compared with 9d, 8Bs and 8Bm (Fig. S5E). On the medial surface of the brain, scattered axon fibers were visible in area F5 (Fig. S5L and M), F7 (Fig. S5L and N), and F2 (Fig. S5L) of the premotor cortex. The axons with the premotor cortex exhibited a gradient pattern with the largest axon distribution along the anterior part (Fig. S5L). In addition, axons were noted in the precentral opercular area (PrCO) and medial prefrontal area (mainly in 10mr) (Fig. 3C). Interestingly, the projections anchored in the prefrontal cortex of these axonal fibers formed isolated clusters (Fig. 4F and H, indicated by arrows). The z-axis extent of axonal clusters was ranging from 1.2 mm to 3.8 mm (2.24 ± 0.80 mm) (Fig. S6).

Beyond the frontal lobe, rich connections were observed in the temporal lobe (Fig. 3), predominantly in caudal lateral (CL), caudal (CPB) and rostral (RPB) portions of parabelt region of the auditory cortex; anterior TE (TEa), medial TE (TEm), superior temporal polysensory area (STP, correspond to areas PGa and TPO), IPa and TAa of the dorsal bank/ventral bank/fundus of the superior temporal sulcus (STSd/v/f) (Fig. 3C); medial superior temporal area (MST), floor of superior temporal area (FST), anteroventral TE (TEav), anterodorsal TE (TEad) and area TEO of superior temporal area. The vlPFC also sends axons to limbic lobe, mainly in 24a, 24a’, 24b, 24b’ and 24c of anterior cingulate areas (ACC) (Fig. 3C); area TF of parahippocampal cortex (Fig. 3C). Relatively weak projections were observed in the dorsal subdivision of lateral intraparietal area (LIPd); 7a and 7b of inferior parietal lobule areas; secondary somatosensory area (SII) and parietal operculum (7op) of the parietal cortex (Fig. 3C). There were some sparsely labelled axons in the granular insula (Ig) and lateral agranular insula (Ial) area (Fig. 3C). In white matter, traveling axonal bundles were found in the corpus callosum, anterior limb of internal capsule (ALIC, Fig. S7A-B), and anchored into the MD thalamus (Fig. S7C-D). Subcortically, axon clusters were observed in the medial (Fig. S7F-G) and caudal (Fig. S7H) parts of caudate. High resolution confocal images revealed that axons in MD (Fig. S7D) and caudate (Fig. S7J) were thinner than those in the ALIC (Fig. S7B). Furthermore, the labelled axons was found extending to the parvicellular part of accessory basal nucleus of amygdala (ABpc), reticulate and compacta parts of substantia nigra (SNr/c), claustrum and subthalamic nucleus (STN) (Fig. 3C).

### Comparison of vlPFC axonal projections by dMRI and STP

We further introduced a quantitative comparison of vlPFC connectivity profile obtained by dMRI-based tractography and STP data. Typical T2-weighted and dMRI images of the macaque brain acquired from an ultra-high field MRI scanner were shown in Fig. 5A-D. To carry out a proof-of-principle investigation, we focused on the vlPFC-CC-contralateral tract that was reconstructed in 3D space by using STP and dMRI data, respectively (Fig. 6A and B). After co-registering the reconstructed tracts into a common 3D space, our approach relied on slice-based statistical correlation methods (the Pearson correlation and Dice coefficients) along this vlPFC-CC-contralateral tract. Upon visual comparison, the dMRI-derived tracts largely overlapped with the axonal bundles shown in STP images (Fig. 6A and B). Statistical correlation indices were computed for each pair of diffusion tractography and STP images to quantify their spatial overlap. We found consistent, marked agreement between these two modalities along this tract (Fig. 6C), as demonstrated in Fig. 6 D-F. For all slices (spaced by 500 μm) along vlPFC-CC-contralateral tract, we observed consistent and significant correlations between these two modalities (R = 0.4368 ± 0.0850; Dice = 0.4061 ± 0.0939). Two example GFP-labelled axon images as marked in Fig. 6F were displayed in Fig. 6G-J with different magnifications, showing typical travelling axons in corpus callosum (Fig. 6 G and H) and frontal white matter (Fig. 6 I and J).

**Figure 5.**
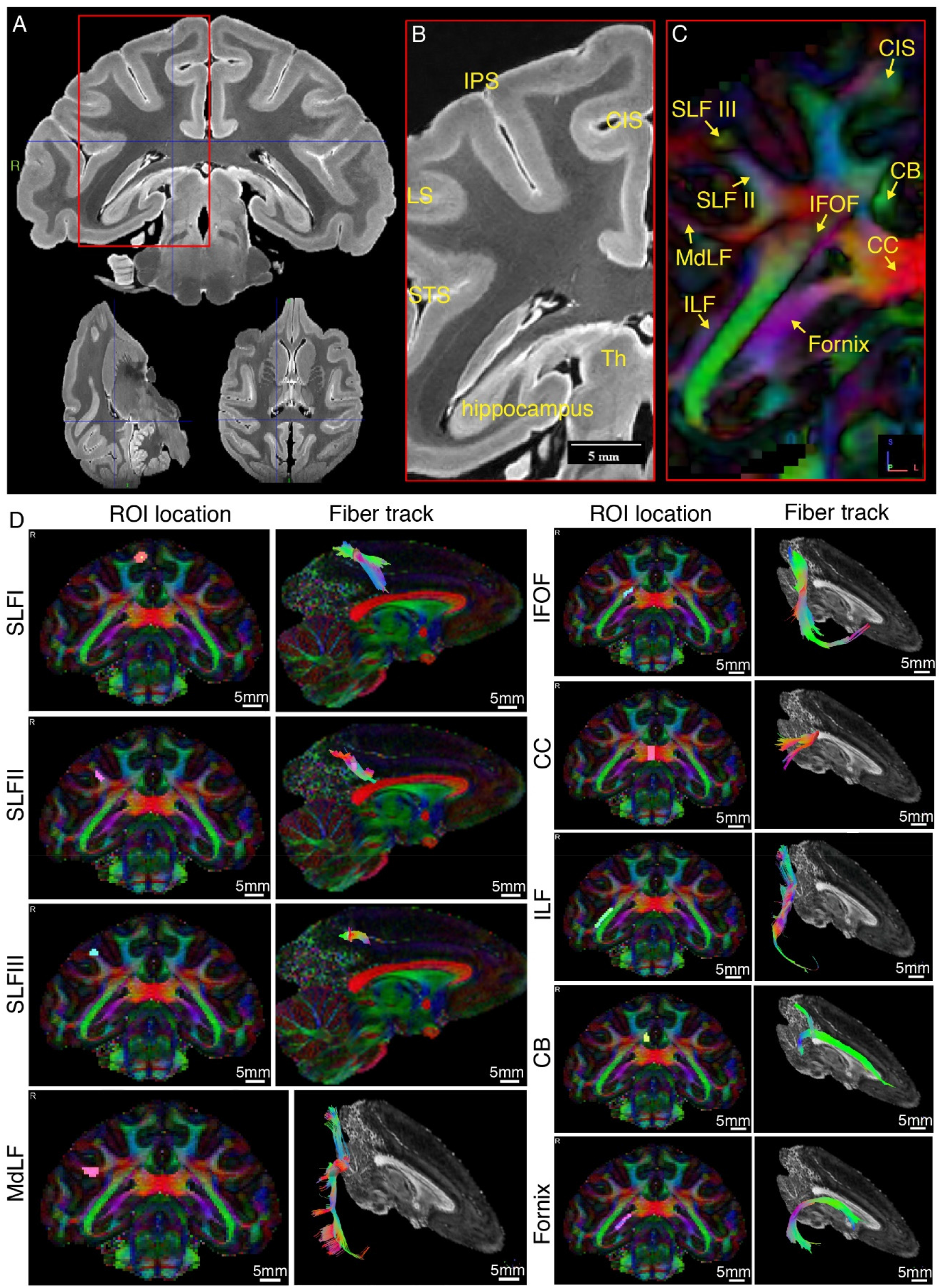
Representative *ex-vivo* MRI images of the macaque brain. (**A**) Typical high-resolution T2-weighted images were shown in axial, coronal, and sagittal planes. (**B**) Zoom-in view of the red box in a, shown with anatomical landmark gyri including intraparietal sulcus (IPS), lunate sulcus (LS), superior temporal sulcus (STS), and cingulate sulcus (CIS). (**C**) The color-coded FA map corresponding to b. Major fiber bundles including superior longitudinal fasciculus subcomponent I, II and III (SLF-I, -II, - III), inferior fronto-occipital fasciculus (IFOF), inferior longitudinal fasciculus (ILF), middle longitudinal fasciculus (MdLF), corpus callosum (CC), cingulum bundle (CB) and fornix are clearly demonstrated. Red color codes left and right, blue color codes anterior and posterior, and green color codes superior and inferior directions. (**D**) Typical tractography of the main fiber bundles indicated in c are derived from the present dMRI data. The ROI locations and fiber tracks are overlaid on the color-coded FA maps.

**Figure 6.**
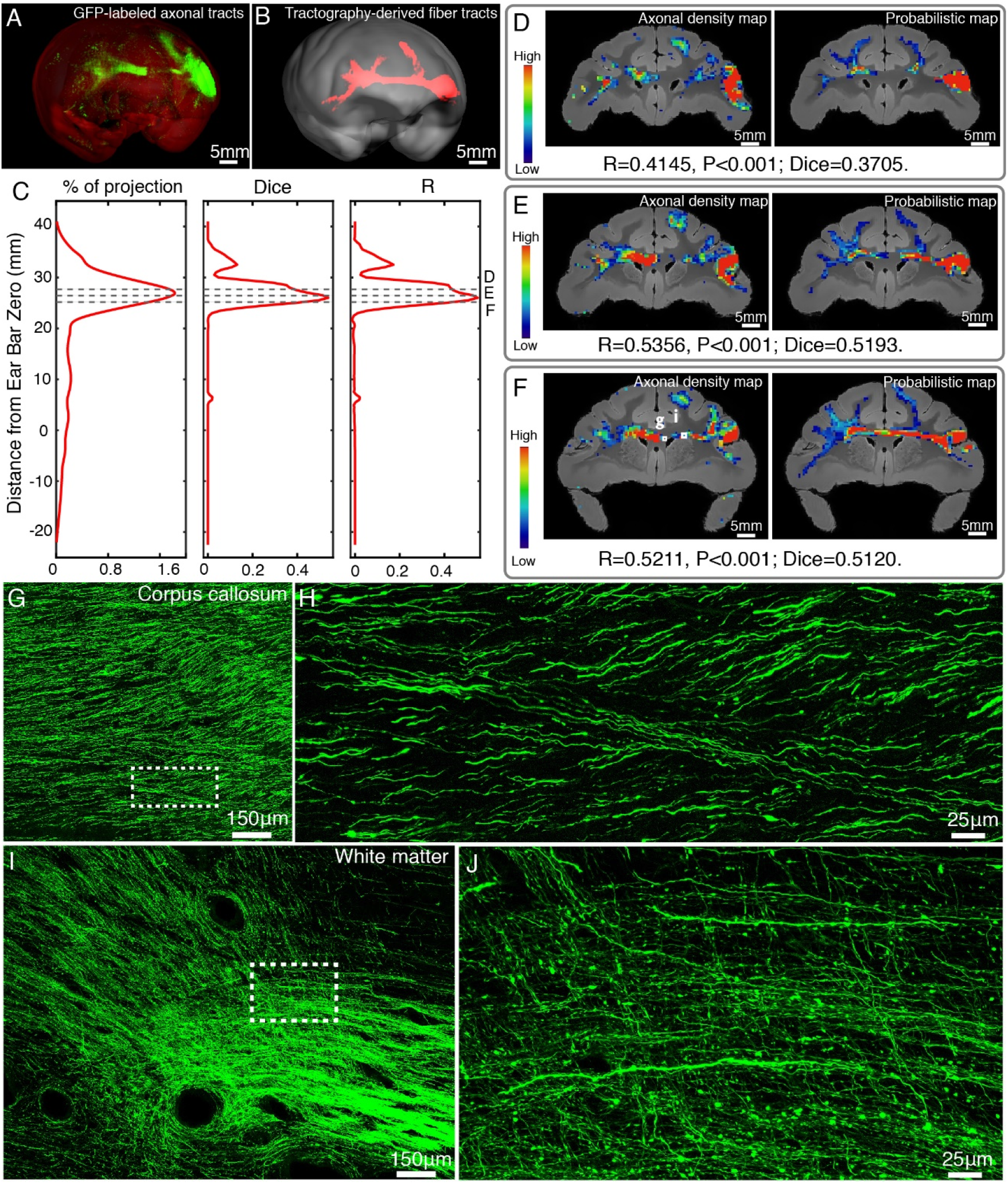
Comparison of vlPFC connectivity profiles by STP tomography and diffusion tractography. (**A**, **B**) 3D visualization of the fiber tracts issued from the injection site in vlPFC to corpus callosum to the contralateral vlPFC by STP tomography and diffusion tractography. (**C**) Percentage of projection, Dice coefficients and Pearson coefficients (R) were plotted along the anterior-posterior axis in the macaque brain. (**D**-**F**) Representative coronal slices of the diffusion tractography map and the axonal density map along the vlPFC-CC-contralateral tract, overlaid with the corresponding anatomical MR images. (**G**-**J**) GFP-labeled axon images as marked in Fig. 6F were shown with magnified views. (H, J) correspond to high magnification images of the white boxes indicated in G and I, both of which presented a great deal of details about axonal morphology.

### Inferior fronto-occipital fasciculus in macaques

As illustrated by diffusion tractography, the inferior fronto-occipital fasciculus (IFOF) in macaques is a long-ranged bowtie-shaped tract (Fig. 7A), showing traveling course similar to humans. The frontal stem of IFOF spread to form a thin sheet, and its temporal stem narrowed in coronal section, mainly gathered at the external capsule. The intersection between IFOF and axonal projections of vlPFC was shown in a common 3D space of diffusion tractography in Fig. 7B, whereby the posterior part of vlPFC axonal projections apparently end at the middle superior temporal region, far from the occipital lobe. To quantify the spatial correspondence between the IFOF tract and vlPFC projectome, the Szymkiewicz-Simpson overlap coefficient was calculated in a shared common 3D space after co-registration. It was 0.038 in 3D space, indicating that only a small fraction of the IFOF tract and vlPFC projectome overlapped (mainly in the front half of the brain, Fig. 7). Also the Szymkiewicz-Simpson overlap coefficients between 2D coronal slices of IFOF and vlPFC projectome was plotted along the anterior-posterior axis of the macaque brain (Fig. 7C). The anterior part of the vlPFC axonal projections shown by STP tomography largely overlapped with the dMRI-derived IFOF tracts in frontal whiter matter (Fig. 7D), external capsule (Fig. 7E and F), claustrum (Fig. 7E and F) and extreme capsule (Fig. 7F). Meanwhile, the posterior part of dMRI-derived IFOF tract passed through temporal white matter (Fig. 7G), whereas the posterior part of fiber projections of vlPFC sent no axons to this region (Fig. 7G).

**Figure 7.**
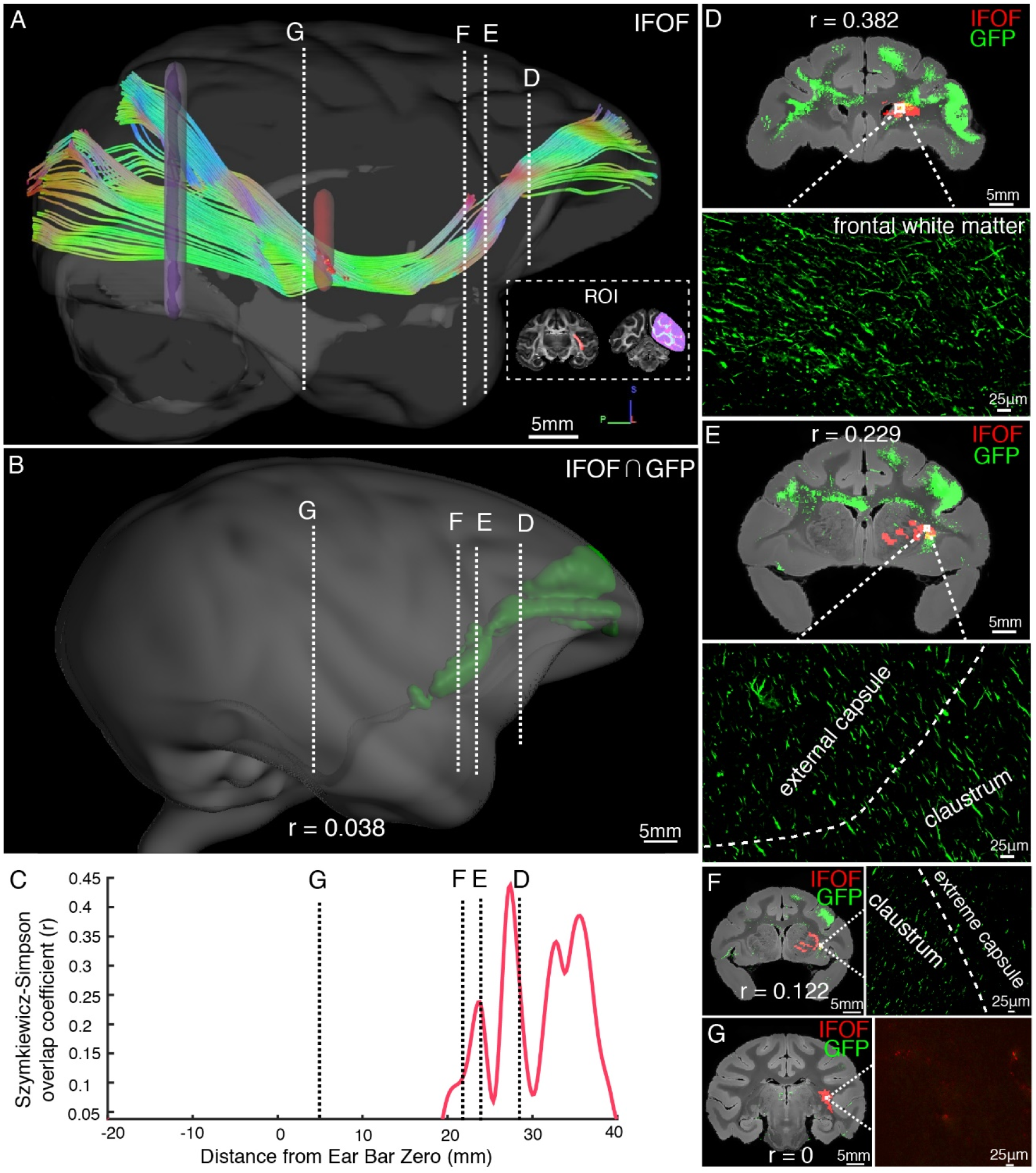
Illustration of the inferior fronto-occipital fasciculus by diffusion tractography and STP. (**A**) The fiber tractography of IFOF (lateral view). Two inclusion ROIs at the external capsule (pink) and the anterior border of the occipital lobe (purple) were used and shown on the coronal plane. The IFOF stems from the frontal lobe, travels along the lateral border of the caudate nucleus and external/extreme capsule, forms a bowtie-like pattern and anchors into the occipital lobe. (**B**) The reconstructed traveling course of IFOF based on vlPFC projectome was shown in 3D space. (**C**) The Szymkiewicz-Simpson overlap coefficients between 2D coronal brain slices of the dMRI-derived IFOF tract and vlPFC projections were plotted along the anterior-posterior axis of the macaque brain. Four cross-sectional slices (D-G) along the IFOF tracts were arbitrarily chosen to demonstrate the spatial correspondence between the diffusion tractography and axonal tracing of STP images. (**D**-**G**) The detected GFP signals (green) of vlPFC projectome and the IFOF tracts (red) obtained by diffusion tractography were overlaid on anatomical MRI images, with a magnified view of the box area. Evidently there was no fluorescent signal detected in the superior temporal area where the dMRI-derived IFOF tract passes through (G).

## Discussion

### Brain-wide excitatory projectome of vlPFC in macaques

We customized STP tomography for whole-brain imaging of the macaque monkey at submicron resolution and accomplished brain-wide 3D reconstruction of axonal connectome, thanks to prominent characteristics of STP tomography including free of tissue distortions, no need for section-to-section alignment, and high-resolution image sets readily warped in 3D space^36^. Importantly, we coupled STP tomography with genetic methods using enhancers/promoter elements that target specific cell types^20^. Previous studies have demonstrated that a CaMKIIα promoter carried by lentivirus was able to target excitatory neurons with optogenetic proteins in the macaque brain^27^, and a TH promoter carried by AAV selectively targeted dopamine neurons^31^. Here we deployed AAV with CaMKIIα promoter to focus on the excitatory projection of vlPFC, whereby immunofluorescent staining with both CaMKIIα and GABA confirmed that GFP was specifically expressed in excitatory neurons. Hence, this integrated approach allows clear dissection of projection patterns from diverse neuronal types^30^, and enriches our knowledge about the anatomical infrastructure of neural circuits for individual cell types at the entire brain scale^31^.

Both anterograde and retrograde tracing evidence shows that vlPFC is extensively connected to other divisions of PFC including OFC, FEF and the ventral premotor cortex. Extrinsic connections beyond the PFC, vlPFC is connected mainly to the dysgranular insula, frontal operculum, somatosensory-related areas in the parietal operculum and inferior parietal cortex, visual-related areas in the inferior temporal cortex, and anterior cingulate areas^6, 37^. We found the excitatory projection of vlPFC to the rest of brain was compatible with previous reports using chemical tracers^3, 37, 38, 39^. Furthermore, we compared the current vlPFC projectome data with the well-known macaque connectivity database CoCoMac^40^, which includes the results of several hundred published axonal tract-tracing studies in the macaque monkey brain^41^. Essentially the vlPFC connectivity profile shown here was markedly similar to that of CoCoMac, except that vlPFC projections to PFCol, PFCdm, PFCdl, PFCoi, PFCm, PMCvl, amygdala, and SII have not been reported in CoCoMac database or reported merely with unspecified strength. In addition, we compared vlPFC projections with one recent report^37^, showing that the brain regions projected from area 45 were clearly observed in the present vlPFC projection data. Note that we used the projection volume index instead of the fluorescence intensity, which has been demonstrated reliably to quantify axonal connectivity strength^4^. Although the passing fiber and terminal were not readily distinguishable, the results of terminal labelling that compared synaptophysin-EGFP-expressing AAV with the cytoplasmic EGFP AAV have shown high correspondence in target areas^4^.

### AAV2/9 is suitable for long-range axonal tracing in the macaque brain

Methods for tissue labeling have been continuously evolving from silver impregnation of degenerating fibers to ex-vivo visualization of axonally-transported tracers injected at single brain nuclei, and finally to an integrated style which coupled high-resolution whole-brain imaging technologies with viral and genetic tracers^28^. Among four viral vectors tested here, we found that AAV2/9 demonstrated the highest efficiency of long-range axonal tracing in the macaque brain. VSV was initially utilized as a transsynaptic tracer in a previous study since VSV encodes five genes, including G protein which promotes anterograde transsynaptic spread among neurons^42^. In our study, we used VSV with G deletions to trace axonal projection without trans-synaptic labeling, which enabled robust gene expression at remarkably higher level relative to other vectors in a very short time (less than a week). But we found that a shorter expression time of VSV-ΔG was insufficient to label axons traveling long distance whereas a longer expression time of VSV-ΔG caused cell death, consistent with a prior finding that VSV-G failed to label transsynaptic cells at distant areas^43^. The advantage of lentivirus, which is derived from human immunodeficiency virus type 1 (HIV-1)^44^, is that it has a large genetic capacity of approximately 10 Kb which allows for the expression of multiple gene and usage of more than one promoter or regulatory elements. And we found GFP expression induced by lentivirus remarkably stable after 9 months in macaque monkeys, even though the labeled level was mild^45^ and the labeled scope was limited.

As an effective carrier for gene delivery into the brain, AAV has a number of established advantages including minimal toxicity, weak host immune response, stable gene expression in neurons with extraordinarily high transfection efficiency (titers up to 10^12^–10^13^ genome copies per mL)^30^. One major drawback of AAV vectors is the limited packaging capacity. AAVs usually deliver gene cassettes of approximately 4.8 Kb (i.e., one or two small genes)^28^, which has motivated us in pursuit of biocompatible nano-based carriers^46^. It is well known that different AAV serotypes have their own sequences in the inverted terminal repeats such that they have distinct transfection bases towards various cell types in the brain. The recombinant virus we used was AAV2/9 which contains the inverted terminal repeats from AAV serotype 2 and the capsid proteins from AAV serotype 9. Previous studies have shown that AAV2 is the most widely-used AAV vector and effectively transfects neurons of non-human primates^47^. In a recent report on a mouse model, researchers co-injected AAV and a classical antegrade tracer - biotinylated dextran amine (BDA) into one brain region and observed long-range projections with similar patterns by both tracers, except that BDA had more retrograde-labelled neurons, probably uptaken by passing fibers in some areas^4^. Together, our results have demonstrated that AAV2/9 vector was more suitable for long-range axonal fiber tracing, while VSV-ΔG was suitable for rapid gene expression and lentivirus for long-term gene expression in macaques.

### Comparison of STP tomography with diffusion tractography

Pioneering studies on cross-modality comparison across the whole-brain scale have been done by constructing a connectivity matrix using dMRI-based tractography and tracer-injection tracing in mice^48^ and in monkeys^18, 49, 50^. The spatial correspondence of axonal fibers derived from diffusion tractography and GFP-labeled fluorescent images have been compared both in mice^51, 52, 53^ and in macaques^54^. Dauguet and coworkers found that the somatosensory and motor tracts derived from diffusion tractography were visually in good agreement with the reconstructed 3D histological sections labeled by anterograde WGA-HRP tracer in a monkey brain, but suffered certain limitations for regions at remote locations from seeds^54^. Moreover, the structural connectivity analyses based on the histological dataset provided varying correlative evidence between these two measurements (like r = 0.21^49^ using the CoCoMac tracer data^40^ and r = 0.59^18^ using the tracer connectivity matrix from^55^). Note that such structural connectivity analysis does not describe a 3D correspondence of the axonal fiber trajectory, but an “end-to-end” match. STP tomography effectively transformed a series of histological slice images into a 3D space with which dMRI-derived tracts were co-registered, thus enabling a direct, quantitative comparison of the high-throughput data from these two modalities. This is technically challenging due to a giant difference in scale between the axonal fibers and image resolution of dMRI^13^. We have taken meticulous steps to maximize the signal-to-noise like using Gd-DTPA as an enhanced contrast agent^56^ and to minimize the image artifacts in an ultrahigh field scanner for achieving a reasonably high spatial resolution. We observed that GFP-labeled axonal density maps not only significantly overlapped with dMRI-derived probabilistic maps throughout the traveling course, but also demonstrated comparable connectivity strengths and patterns. But caution should be born in mind that diffusion tractography estimated the Brownian motion of water molecules, from which the directionality of axons cannot be distinguished^17^. The viral tracing data here contained only anterograde axonal fiber projections.

Our particular focus on the vlPFC connectivity profile leads us to clarify the existence of the IFOF in monkeys which is heavily debated^35^. The IFOF in human brain was first described in the early twentieth century^57^, whereby the anatomy of this pathway in human has been recently shown by micro-dissection and diffusion tractography studies^58, 59^. Its entire course through the ventral part of the external capsule (EC) connects the occipital cortex and the parietal and temporal cortices to the frontal cortex^60^. Some axonal tracing studies showed connections between frontal and occipital lobes in monkeys^37, 55^, which was consistent with the observation by tractography^61^ and blunt dissection^60^ experiments. By contrast, other studies that are capable of tracking monosynaptic pathways failed^62^. Using the same ROIs seeds as prior studies^61, 63^, our ex-vivo tractography did show fiber connections between frontal and occipital lobes in monkeys, matching the trajectory of IFOF in humans. By contrast, using anterograde AAV vector without trans-synaptic capability, we found that vlPFC fiber projections passed through external capsule, claustrum and extreme capsule and anchored to the middle superior temporal region. Although the trajectory of vlPFC between frontal and temporal regions matched well with the diffusion tractography of IFOF, axonal projections of vlPFC never reached the occipital lobe. Lack of monosynaptic tracing data in human subjects, we could not rule out the possibility of same scenario for IFOF in humans. If the IFOF connects the frontal lobe with the occipital lobe in a trans-synaptic manner, it unveils a hitherto unknown information relay/integration process occurring in superior temporal area of the primate species which holds great implications for neural network computation. Nevertheless, unlike the direct monosynaptic connections reported between subdivisions of PFC such as OFC and the visual cortex in mice ^5, 64^, our results underscore a nontrivial species difference and raise interesting questions about the long-range brain organization and the functional role of superior temporal area in primates which definitely merits future examination.

In summary, we present a detailed excitatory connectivity projection map from vlPFC to the entire macaque brain, and demonstrate a broadly applicable roadmap of integrating 3D STP tomography labeled with antero-/retro-grade tracer and diffusion tractography for the mesoscopic mapping of brain circuits in the primate species.

## Materials and Methods

### Animals and Ethics statement

All experimental procedures for nonhuman primate research in this study were approved by the Animal Care Committee of Shanghai Institutes for Biological Science, Chinese Academy of Science, and conformed to the National Institutes of Health guidelines for the humane care and use of laboratory animals. From November 2015 till November 2019, ten adult macaque monkeys (Macaca mulatta and Macaca fascicularis) weighting 3.5 to 12.2 kg (7.0 ± 2.9 kg) were used for in this study (Table 1), two of which (Macaca fascicularis) were used for ex-vivo ultrahigh field dMRI scanning.

**Table 1.**
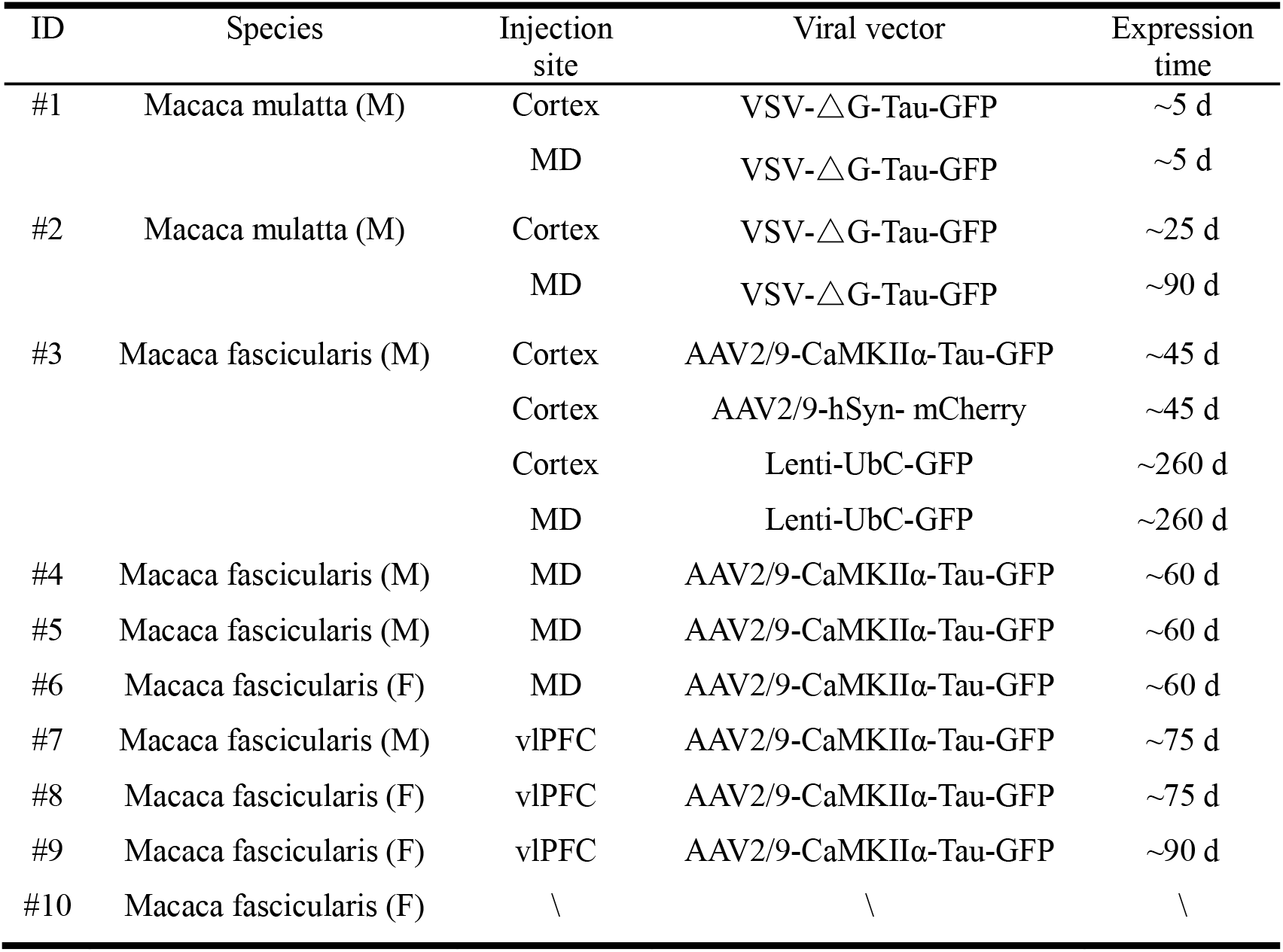
Injection cases and viral vectors used in this study. Abbreviations: M, male; F, female. MD, mediodorsal thalamus; vlPFC, ventrolateral prefrontal cortex; UbC, human ubiquitin C; hSyn, human synapsin I; CaMKII, Ca2+/calmodulin dependent protein kinase II; d, day.

### Viral vectors

Four viral vectors, including VSV-ΔG (VSV-ΔG-Tau-GFP, titer: 5.0×10^8^ PFU/mL), lentivirus (lentivirus-UbC-GFP, titer: 1.33×10^9^ TU/mL), and two constructs of AAV2/9 (AAV2/9-CaMKIIα-Tau-GFP, titer: 8.47×10^13^ vg/mL; AAV2/9-hSyn-mCherry, titer: 1.6×10^13^ vg/mL), were tested in this study (Table. 1). AAV2/9 and VSV-ΔG were purchased from BrainVTA technology Co., Ltd. (Wuhan, China), and lentivirus was provided by a coauthor Z.Q’s laboratory. Here, the recombinant AAV2/9 contained either a hSyn or CaMKIIα promoter to regulate the expression of either reporter gene mCherry in all neurons or fused Tau-GFP protein in glutamatergic excitatory neurons, respectively. Regarding the VSV vector, G protein was deleted to prevent transsynaptic spread. The last tested viral vector, Ubic promoter-driven lentivirus, expressed GFP in all eukaryotic cells.

### MRI-guided virus injection, histology and microscopy

To precisely target brain regions in individual subjects, we performed in-vivo MRI scanning in monkeys ^9, 14, 65, 66^ and then used MRI images to guide the virus injection (more details in Supplementary materials). According to the expression time of individual virus (Table 1), animals were deeply anesthetized, and then transcardially perfused with 0.9% NaCl (pH = 7.2) followed by ice-cold 4% paraformaldehyde in 0.01 M phosphate buffered saline. Brains were extracted and post-fixed in 4% PFA for 3 days. Cryo-sectioning combined with wide field microscope imaging and confocal laser microscope imaging was performed for virus testing and more details were provided in Supplementary materials. Fluorescence signals of AAV labeled areas were detected and recorded using a customized STP tomography (Fig. S8, Supplementary materials). High x-y resolution (0.95 μm/pixel) serial 2D images were acquired at a z-interval of 200 μm across the entire macaque brain, as resulted in a continuous ∼1 month scanning and ∼5 TB STP tomography data for one monkey brain (Fig. S9). Once finished scanning, all sections were retrieved from the cutting bath and stored in cryo protection solution (containing 30% glycol, 30% sucrose in PBS) at −20°C for further histological examination.

### Fluorescence image preprocessing

Fluorescent images of the macaque brain usually contain strong autofluorescence signal (Fig. S10 A-E), mainly caused by the accumulation of lipofuscin^67^. Autofluorescence provides good contrast between gray matter and white matter, which is rather useful for image registration. But the presence of autofluorescence is undesirable for the axon tracing procedure since this background signal sometimes is much stronger than that of some thin GFP labeled axons (Fig. S10 A-E). Nevertheless, thanks to the broad emission spectrum of lipofuscin^68^, autofluorescence and GFP signals are easily distinguishable from each other. We therefore implemented and compared the following three methods for background reduction: (1) transforming the GFP signal from the green channel (488 nm) to the blue channel (405 nm) using immunofluorescent staining (Fig. S10M), (2) subtracting the normalized autofluorescence signal in the red channel from the green channel (Fig. S10F), which contains both GFP signal and autofluorescence background signal, (3) supervised machine learning for autofluorescence exclusion (Fig. S10J).

The first method involved staining the brain tissue with anti-GFP antibody and Alexa Fluor 405 conjugated secondary antibody to transform the GFP signal from green channel to blue channel. Unlike the green and red channels, the transferred blue channel (Fig. S10M) did not contain high intensity autofluorescence puncta. Although this post-hoc thick-section immunofluorescent method successfully reduced autofluorescence, it was incompatible with the block face imaging method. The second one was to subtract the normalized red channel from the green channel using the broad emission spectrum characteristic of autofluorescence puncta, which was able to remove high intensity background signal (Fig. S10F). The third was based on a supervised machine learning plugin for ImageJ, trainable WEKA segmentation^69^, which classifies and binarizes GFP and autofluorescence background signal for background exclusion (Fig. S10J). Both subtraction and machine learning methods were used for better visualization of fluorescence images when necessary, whereas only the supervised machine learning approach was used for quantitative analysis of STP data^68^.

### STP image processing

STP tomography data processing included axonal fiber detection, image stitching, down sampling, cross-modality registration and quantification. The GFP labelled axonal fibers were segmented by using a machine learning algorithm to remove background. During STP tomography scanning, each field of view (FOV) was saved as a 1024 × 1024-pixel image. For image stitching, individual FOV images from red channel, green channel and segmented GFP signal were stitched into full tissue sections using the Terastitcher software. A convolutional neural network-based denoising approach was used to improve SNR of images when necessary^70^. The natural alignment of serial images generated by STP tomography allowed to stack the section images to form a coherent reconstructed 3D volume ^33^. In order to localize the virus injection site, a threshold was set at green channel to retain the fluorescence signal only from the cell soma for each section image. Images of the red channel and injection site volumes were downsampled to a resolution of 200 × 200 × 200 μm grid. For serial segmented GFP images, the total signal intensity was computed for each 200 × 200 μm grid by summing the number of signal-positive pixels in that voxel. Red channel volume was used to perform registration to the monkey brain template, as red channel images contain visible anatomical information of brain structures^71^. The brain template of cynomolgus macaque was adopted from an MRI-based atlas generated from 162 cynomolgus monkeys^66^. We warped the red channel volume to the template space by using a symmetric normalization (SyN) algorithm in ANTs (Fig. S11). The cortical label was adopted from the D99 parcellation map^72^, and subcortical label was adopted from INIA19 parcellation map^73^. Also the segmented GFP volume and injection site volume were co-registered onto the same template. Density of GFP signal and total GFP volume in each parcellated brain region were used to represent the axonal connectivity strength. Percent of total projection was defined by the GFP-positive pixel count within each parcellated brain region (or brain lobe) normalized to the total of all GFP-positive pixels. Additionally, the percent innervation density was calculated as the proportion of density of GFP pixel counts covering the maximal density of GFP pixel counts of the brain. To create plots that display the data along the anterior-posterior axis (e.g. % density innervation), the location of ear bar zero was used as the origin. The percent innervation density of each cortical region innervated by vlPFC was rendered onto a brain surface.

### Ex-vivo MRI scanning and data preprocessing

We collected dMRI data using an 11.7 T horizontal MRI system (Bruker Biospec 117/16 USR, Ettlingen, Germany), equipped with a 72 mm volume resonator and an actively shielded, high performance BGA-S series gradient system (gradient strength: 740 mT/m, slew rate: 6660T/m/s). The paraformaldehyde-fixed macaque brain has immersed in gadolinium MR contrast agent (Magnevist®, Bayer Pharma AG, Germany) mixed with phosphate buffered saline (PBS) solution two weeks before MRI scanning. Then the brain was positioned securely in the close-fitting designed container filled with FOMBLIN® perfluoropolyether (Solvay Solexis Inc. Thorofare, NJ, USA). Air bubbles were removed with vacuum pump for 24 h before scanning. Diffusion MRI data were acquired using a 3D diffusion weighted spin echo pulse sequence with single-line read-out, containing 60 diffusion gradient directions (b = 4000 s/mm^2^) and 5 non-diffusion-weighted (b = 0 s/mm^2^). The scanning parameters were set: TR/TE = 82/22.19 ms, FOV = 64×54 mm, acquisition matrix = 128×108, slice thickness = 0.5 mm, and averages = 3. In addition, whole brain T1-weighted and T2-weighted structural images were obtained using 3D FLASH and 2D Turbo RARE sequences, respectively. All scanning was performed at room temperature (approximately 20 °C) and the total scan time was approximately 36 hours. The dMRI data was preprocessed using the FSL software (http://www.fmrib.ox.ac.uk/fsl)^74^. Individual image volumes were co-registered with b = 0 images to account for eddy currents and B0 drift using affine registration in FLIRT^75^. A custom in-house script was applied to reorient the corresponding gradient direction matrix. Careful steps have been taken to minimize artifacts caused by motion and field distortion, and image correction was applied only if necessary^76, 77^. More details of data acquisition and preprocessing were described in Supplementary materials.

### Reconstruction and comparison of diffusion tractography and axonal tracing

We first identified the injection-site volume in the vlPFC in STP tomography data, warped it to the space of dMRI volume and used it as a seed mask for tractography. Then the injection site related tractography was constructed using the preprocessed dMRI images in FSL toolbox. BEDPOSTX was used for Bayesian estimation of a crossing fiber model with three-fiber orientation structure for each voxel using Markov chain-Monte Carlo sampling^74^. This provided a voxel-wise estimate of the angular distribution of local tract direction for each fiber, which was a starting point for tractography. Tractography was then performed from the injection-site seed masks without waypoint mask and termination mask using the Probtrackx probabilistic tractography software^74^. A probabilistic map of fiber tracts was generated with 500 μm isotropic resolution. A probabilistic map provided, at each voxel, a connectivity value, corresponding to the total number of samples that passed from the seed region through that voxel. The following settings were used: number of samples per voxel = 5000, number of steps per sample = 2000, step length = 0.2 mm, loop check, default curvature threshold = 0.2 (corresponding to a minimum angle of approximately ±80 degrees), subsidiary fiber volume threshold = 0.01, seed sphere sampling = 0 and no way-point or termination mask. In the resulting map, each voxel’s value represented the degree of connectivity between it and the seed voxels. To generate dMRI-derived fiber tracts, the resulting probabilistic maps were set at 0.2% threshold of total number, i.e., any voxel below threshold was set to zero. In parallel, segmented fluorescence images from the STP tomography data were downsampled to 500 μm grid to generate axonal density maps. The signal intensity of an axonal density map was computed for each 500 × 500 μm^2^ grid by summing the number of GFP-positive pixels within that area. Note that the axonal density map was also filtered by setting an intensity threshold of 10^3.2^ to minimize false positives due to segmentation artefacts^4^. After co-registered the probabilistic maps and the axonal density maps onto the same template, both Dice coefficients and pixel-wise Pearson coefficients were calculated to quantitatively assess the spatial overlap^78, 79^.

As described recently^63^, the inferior fronto-occipital fasciculus was reconstructed using streamline-based probabilistic tractography. We ran this probabilistic tractography tool in MRTrix3 (www.mrtrix.org) via bootstrapping^80^. Streamlines were seeded over the whole brain area that encapsulated the tract of interest. Two inclusion masks were used to define two regions that each tract must pass through, and only streamlines that pass through both regions are retained. One exclusion mask was used to restrict tracking to the contralateral hemisphere of the brain. The inclusion and exclusion masks were drawn manually as described previously^63^: the first mask was placed on the anterior border of the occipital lobe in the coronal view, the second mask was placed on the external/extreme capsules in the coronal view and the third mask was cover the whole left hemisphere as the exclusion mask. The delineation process was performed using the MRIcro (www.mricro.com) software. Using this “waypoint” method, the resultant streamlines were able to meet our preset conditions. More details of streamline-based probabilistic tractography processing were described in Supplementary materials. To further reduce false positive tracts, any streamlines that were identified as either attached to other tracts or anatomically implausible trajectories were manually removed. For a direct comparison between diffusion-derived the IFOF tract and vlPFC projection fibers, we first generated the track density images of the IFOF tract and co-registered them onto the space of the template. The spatial overlap of GFP-positive vlPFC projection fibers and the IFOF tract were then detected with using ImageJ and FSL software in both 2D and 3D space (Fig. 7B). The Szymkiewicz-Simpson overlap coefficient was adopted to quantify the spatial relationship between the IFOF tract and vlPFC projectome, which was defined as the size of the union of them over the size of the smaller set:

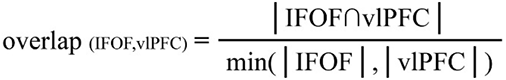

The Szymkiewicz-Simpson overlap coefficient ranges from 0 (no overlap) to 1 (if the IFOF tract is found in its entirety in vlPFC projectome).

### Materials availability

All the virus vectors used in this paper are available from the authors for sample test.

### Data Availability

There is no institutional resource for hosting such big connectome data. Therefore we host it ourselves on a publicly accessible FTP secure server, ftp://192.168.220.53 via VPN https://119.78.67.35:666, login: ftpuser1, password: ftp@user1. We commit to keeping it available for at least 5 years, and provide alternative procedures where users can copy any or all of it to their own computer.

## Acknowledgments

We would like to thank Jinqiang Peng and Jie Xu for their assistance to data acquisition, and thank Drs. John Gore, Ed Callaway and Anna Roe for their stimulating discussions and suggestions during the preparation of this study. This work was supported by the Key-Area Research and Development Program of Guangdong Province (2019B030335001), National Key R&D Program of China (No. 2017YFC1310400; No. 2018YFC1313803), the Strategic Priority Research Program of Chinese Academy of Science (No. XDB32000000), grants from National Natural Science Foundation (31771174), and Shanghai Municipal Science and Technology Major Project (No. 2018SHZDZX05).

## Author Contributions

Y.M.C., L.Q., L.Q.M., B.T.T., C.X.Y., L.Y.L., Z.Y.F., and Y.S.Y. participated in-vivo MRI scanning and virus injection experiments. Y.M.C. with the help of L.Q., L.Q.M., and S.X.Y. performed immunohistology and STP experiments and processed data. Y.W.W. performed ex-vivo MRI experiment with the help of Y.B.F., Z.X.Y., W.H., and processed data with the help of L.Q., L.Q.M., and F.Y.J. X.F.Q. and Q.Z.L. designed and prepared the viral vectors. W.Z., Y.M.C. and X.F.Q. designed the study and W.Z. supervised the study. Y.M.C., Y.W.W. and W.Z. prepared the manuscript with input from all of the authors.

## Supplementary Information for

### This PDF file includes

Supplementary text

Figures S1 to S11

References

### Supplementary Methods

#### MRI-guided virus injection

T1 weighted images for each monkey were obtained with a 3T scanner (Siemens Tim Trio, Erlangen, Germany) under general anesthesia. A detailed description of in-vivo MRI scanning procedure has been described in our previous studies^1, 2, 3, 4, 5, 6^ and briefly summarized here. Anesthesia was induced by intramuscular injection of ketamine (10 mg per kg). Deep anesthesia was maintained by isoflurane (1.5-3%) and vital physiological signals were continuously monitored during MRI scanning. Anatomical scans were acquired with an MPRAGE sequence using the following parameters: TR = 2300 ms, TE = 2.8 ms, TI = 1100 ms, spatial resolution 0.5 mm isotropic. The target regions were localized in each animal by warping the 3D digital atlas of Saleem and Logothetis^7^ to the individual T1 image using a symmetric normalization (SyN) algorithm. The location of the vlPFC was then calculated with regard to the stereotaxic space.

All procedures for virus injection were performed in strict aseptic conditions. The head of the animal was fixed in a stereotaxic apparatus, within the same coordinate space as the MRI images. The target area was then labelled and an incision was made to expose the skull. A burr hole with a 2 mm radius was drilled above the target according to the calculated coordinates, and the dura was carefully incised to expose the cortical surface. The viral vector was delivered into the cortex using a 33-gauge Hamilton syringe controlled by an UltraMicroPump and a micro4 controller (WPI). The injection speed started with 200 nl/min and was increased to 400 nl/min; total injection volume was 10-20 μl. After injection, the needle was retained for at least 15 minutes and drawn back at a rate of ∼1 mm/min. The burr hole was then filled with bone wax and the skin was sutured. Cephalosporin was given for three consecutive days after surgery (25 mg/kg/day, i.m., once a day).

#### Cryo-sectioning

For virus testing, serial sections were cut on a freezing microtome. The fixed brain was first cut into a block, then equilibrated sequentially in 15% and 30% sucrose in PBS until it sank to the bottom of the container. A cryostat microtome (Leica CM1950) was used to serially slice the brain into 50 μm sections. Brain slices were preserved in a cryoprotectant solution (containing 30% ethanediol, 30% sucrose in PBS solution, pH = 7.2) for further immunofluorescence staining and imaging.

#### Serial two-photon tomography

To image the monkey brain, we customized the STP tomography system which was integrated a two-photon microscope (Bruker) with a vibratome (WPI) (Fig. S8), computer controlled and fully automated. The XY stage covered a 50*60 mm^2^ area, and the 3D scanning of Z-volume stacks was achieved with using a stepper motor (Thorlabs) that traveled over 70 mm. The fixed brain that was embedded with 4% agarose was scanned in a 3T MRI to obtain ex-vivo T1 images. Using these T1 images as reference, the active imaged region of each section was determined during STP tomography for improved imaging efficiency. The embedded brain was then held via a magnetic adaptor to a stepper motor and immersed in a cutting bath filled with PBS containing 0.1% sodium azide. The vibratome blade was aligned in parallel with the leading edge of the specimen block. Brain images were captured from the anterior PFC to posterior V1 in the coronal plane. Fluorescence signals for the green channel (excitation wavelength light in 920 nm) and red channel (excitation wavelength light in 1045 nm) were acquired at 30 μm below the cutting surface through a Nikon 16x Water objective (N.A. = 0.8).

During serial scanning, the STP system was fully automated: each optical section was imaged as a mosaic of fields of view on the block surface as the xy stage moved the brain under the objective; once an entire section was imaged, the xy stage moved the brain to the vibratome and cut off a 200 μm section from the top of the sample. The remaining specimen was then moved back under the objective for imaging the next neighboring plane. Optical and mechanical sectioning were repeated until the complete brain data was collected. Hence fluorescent images of the whole monkey brain were continuously acquired (Fig. S9).

#### Histological staining

To perform immunofluorescence procedure, brain slices were incubated in blocking solution containing 5% BSA and 0.3% Triton X-100 in PBS at room temperature for 2 hr and then overnight with primary antibodies in PBS containing 3% BSA and 0.3% Triton X-100 at 4 °C. Slices were rinsed in PBS followed by Alexa Fluor-conjugated secondary antibodies at room temperature for 3 hrs, and DAPI (Cell signaling Cat# 4083s) for 30 mins at room temperature. The following primary antibodies were used: CaMKIIa (1:200, Abcam, Cat# ab5683, RRID: AB_305050), GABA (1:200, Abcam Cat# ab8891, RRID: AB_306844), NeuN (1:500, Millipore, Cat# MAB377, RRID: AB_2298772), GFP (1:300, Thermo Fisher Scientific, Cat# A-11122, RRID: AB_221569). The following secondary antibodies were used: Goat anti-Rabbit IgG (H+L) Alexa Fluor 405 (1:500, Thermo Fisher Scientific, Cat# A31556, RRID: AB_221605), Donkey anti-Rabbit IgG (H+L) Alexa Fluor 568 (1:600, Thermo Fisher Scientific, Cat# A10042, RRID: AB_2534017), Goat anti-Mouse IgG (H+L) Alexa Fluor Plus 647 (1:500, Thermo Fisher Scientific Cat# A32728 RRID: AB_2633277), Goat anti-Rabbit IgG (H+L) Alexa Fluor 488 (1:300, Thermo Fisher Scientific, Cat# A11034, RRID: AB_2576217). The brain slices were mounted onto customized 2 × 3 inch or 3 × 4 inch glass slides. The sections were then scanned using an Olympus VS120 (Olympus, Japan), a wide field microscope, with a U Plan Super Apo 10× objective (N.A. = 0.4) at a resolution of 0.65 μm/pixel. High resolution fluorescent images were acquired with a confocal laser microscope Nikon TiE (Nikon, Tokyo, Japan) with a Plan Fluo 40× Oil DIC N2 objective (N.A. = 1.3), 0.5 μm Z-interval, and 1024 × 1024 pixels.

#### Ex-vivo diffusion MRI

After a fixation period of ∼ 30 days, the whole brain specimen was immersed in a 1:100 dilution of a 1 mmol/mL gadolinium MR contrast agent (Magnevist®, Bayer Pharma AG, Germany) mixed with phosphate buffered saline (PBS) solution for 14 days. Before MRI scanning, the specimen was washed and drained of water from the surface, then positioned into a customized container which was 3D printed for perfect accommodation of the brain sample. Thus the brain was held steadily during MRI scanning. And the container was filled with FOMBLIN® perfluoropolyether (Solvay Solexis Inc. Thorofare, NJ, USA) for susceptibility matching and improved magnetic field homogeneity. The specimen was degassed with a vacuum pump for 24h under 0.1 atmosphere pressure to remove all air bubbles in the sample at 20 °C (magnet room temperature). The ex-vivo macaque brain was scanned on a 11.7 T animal MRI system (Bruker Biospec 117/16 USR, Ettlingen, Germany), equipped with a 72 mm volume resonator and an actively shielded, high performance BGA-S series gradient system (gradient strength:740 mT/m, slew rate: 6660T/m/s). dMRI images were acquired using a 3D diffusion-weighted spin echo pulse sequence with single-line read-out, TR/TE = 82/22.19 ms, FOV = 64×54 mm, matrix = 128×108, slice thickness = 0.5 mm and averages = 3, which included 60 diffusion directions with b = 4000 s/mm^2^ (Δ/δ = 15/2.8 ms, maximum b value = 4234.97, gradient amplitude = 97.19 mT/m) and five non-diffusion encoding (b = 0 s/mm^2^) directions. For the ex-vivo diffusion MRI data acquisition, the *b*-value was recommended to set at 4000 s/mm^2^ 8, 9. T2 weighted images were acquired using a 2D Turbo RARE sequence with TR/TE = 8353.42/28.8 ms, flip angle = 87°, matrix = 450×450, FOV = 54×45 mm, slice thickness = 0.5mm, and averages = 6. T1 weighted images were acquired using 3D FLASH sequence with TR/TE = 40/5.5 ms, flip angle = 15°, matrix = 290×225, FOV =58×45 mm, slice thickness = 0.2 mm, and averages = 4. The total scan time was approximately 36 hours.

Visual inspection of MRI data was first performed to ensure that there were no obvious image artefacts and geometric distortions. Then we calculated the signal-to-noise ratio (SNR) for typical diffusion images. As diffusion images were acquired by spin warp imaging (image reconstruction by a 3D Fourier transform) with a volume quadrature coil, the SNRs were calculated using the “two-region” approach^10, 11^. Specifically, for each gradient encoding direction, the deep white matter (WM) were extracted in subject-native diffusion space to represent the signal^12^; a region positioned in the no signal area at the corner of the image was used to represent the noise. As a rule of thumb, the SNR of b = 0 s/mm^2^ images should be minimally larger than 20 for obtaining relatively unbiased measures of parameters such as FA^13^. Typical SNRs of diffusion images with b = 0 and b = 4000 in the present study were 48.34 ± 8.50 and 23.13 ± 2.05, respectively. It allowed a reliable seed-based 3D reconstruction for diffusion tractography, as illustrated in Fig. 5.

#### Construction of inferior fronto-occipital fasciculus

The streamline-based probabilistic tractography strategy was used to generate the IFOF tracts in 3D^14^. The fiber orientation distribution function (FOD) was estimated with MRtrix3 (www.mrtrix.org)^15^ using the *tournier* algorithm for single-tissue Constrained spherical deconvolution^16^. For fiber tracking, we then used *tckgen* with the *Tensor_Prob* tracking algorithm in MRtrix3^17^. Within each image voxel, a residual bootstrap was performed to obtain a unique realisation of the dMRI data in that voxel for each streamline. These data are then sampled via trilinear interpolation at each streamline step, the diffusion tensor model is fitted, and the streamline follows the orientation of the principal eigenvector of that tensor. The following additional tckgen settings and inputs were used: step size of 0.25 mm, max. angle between successive steps = 45°, max. length = 150 mm, min. length value set the min. length 10 mm, cutoff FA value = 0.1, b-vectors and b-values from the diffusion-weighted gradient scheme in the FSL format, b-value scaling mode = true, maximum number of fibers = 10,000, and unidirectional tracking.

---

**Fig. S1.**
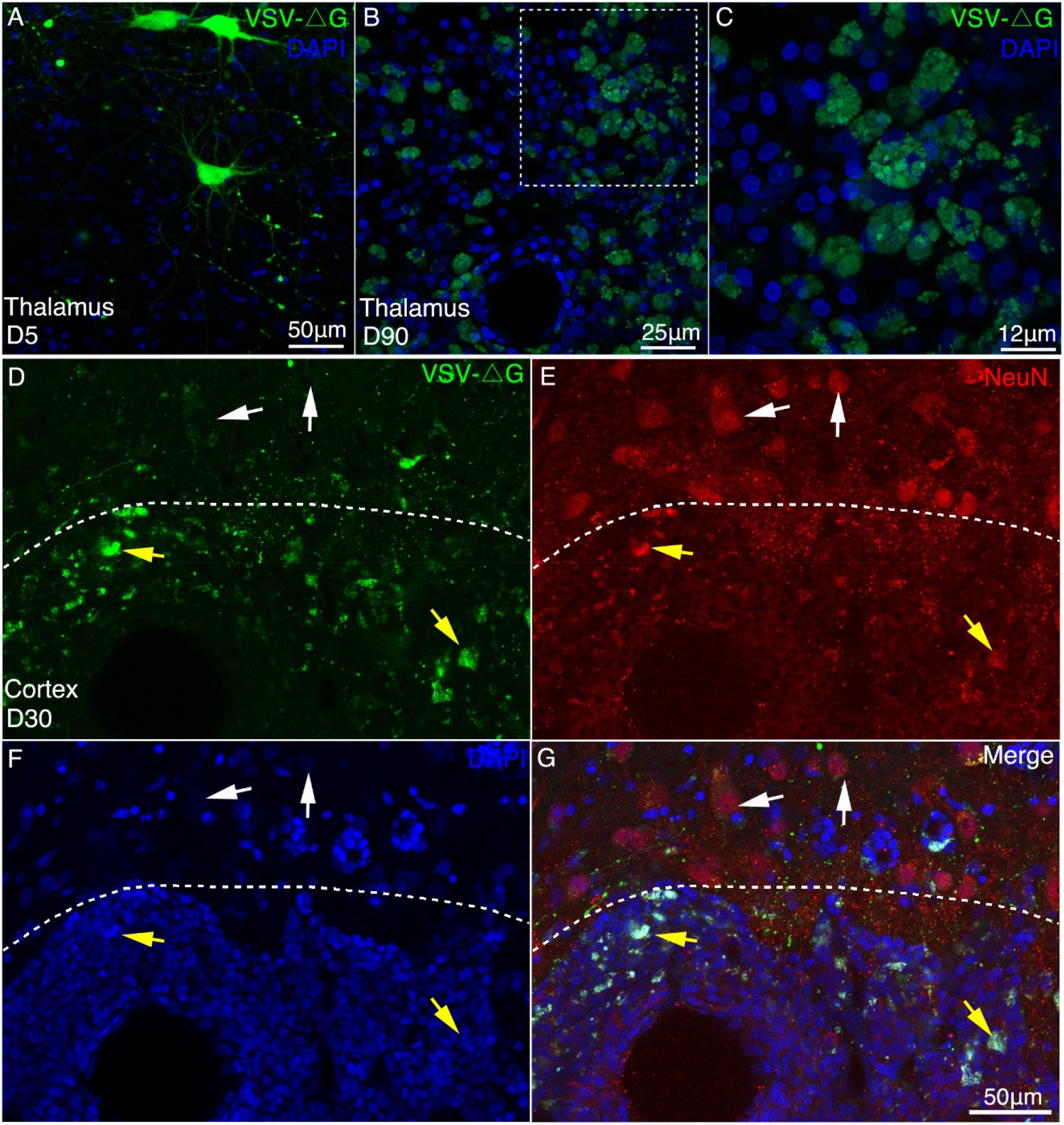
Long-term expression of VSV-ΔG induced neurotoxicity in macaque brain. (**A**-**C**) Compared with the short-term expression (A, 5 days), severe morphological abnormalities such as neurites loss and membrane blebbing (C), were induced by VSV (ΔG) after long-term expression (90 days). (**D**-**G**) 30 days after injection of VSV, immunofluorescent staining was performed with NeuN antibody in the injection site (below the dotted line) and non-injection site (above the dotted line). The non-infected neurons were found to be EGFP negative and neuronal-specific nuclear protein (NeuN) positive by immunofluorescent examination. In the injection site, VSV-infected neurons with GFP and NeuN positive exhibited apparent morphological abnormalities.

**Fig. S2.**
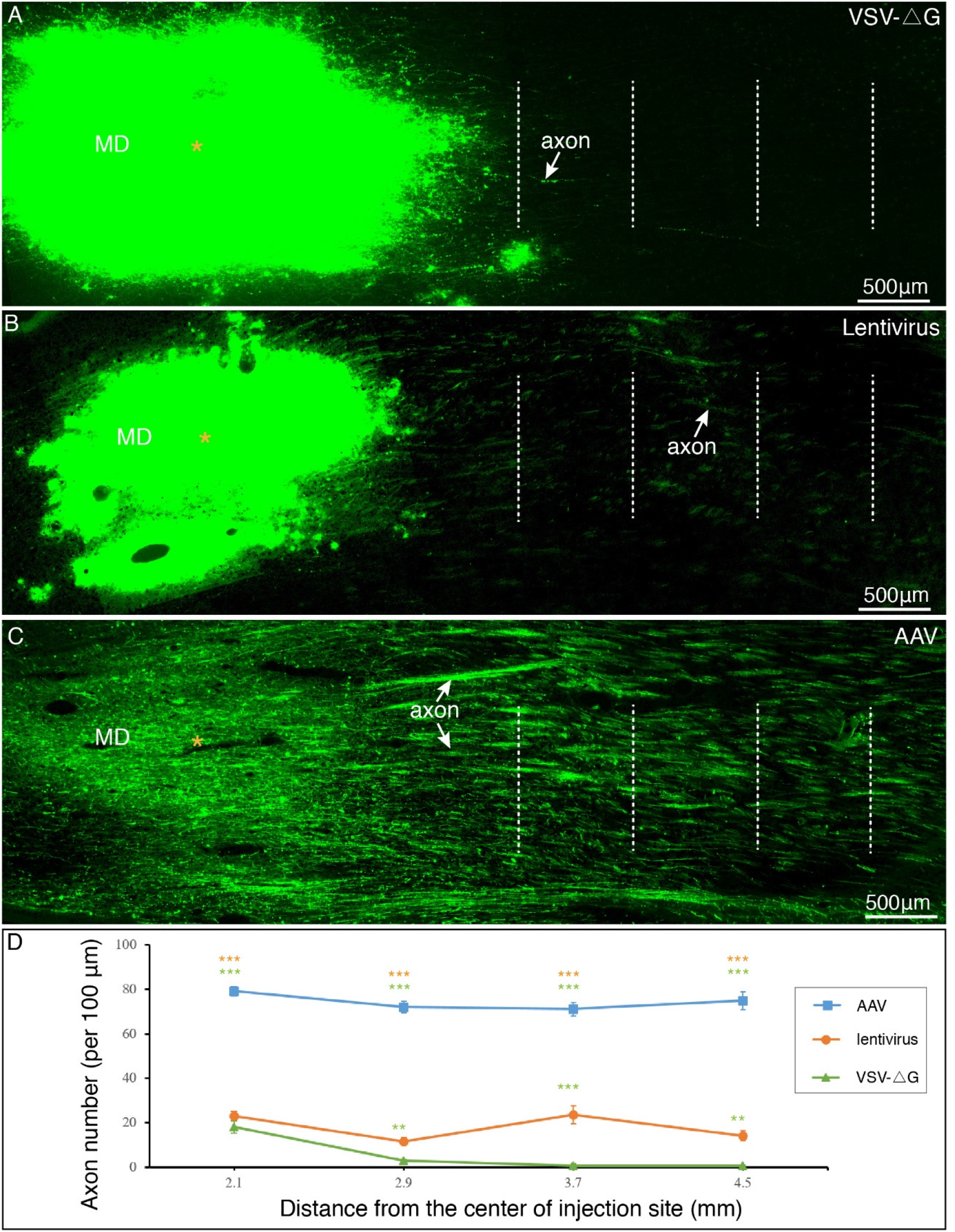
Long-range axonal tracing outcomes using AAV2/9, lentivirus and VSV-ΔG. (**A**) VSV rarely labeled long-range axonal fibers issued from MD thalamus. (**B**) Only sparse axon fibers projecting from MD thalamus were labeled with using Lentivirus. (**C**) Robust projecting fibers were detected distant from the MD thalamus with using AAV2/9. (**D**) Quantitative comparisons among three viral vectors. The number of GFP-labeled axons was counted at each dotted line in a-c, and subject to One-way ANOVA followed by Bonferroni Correction. **p < 0.01; ***p < 0.001; error bars represent SEM.

**Fig. S3.**
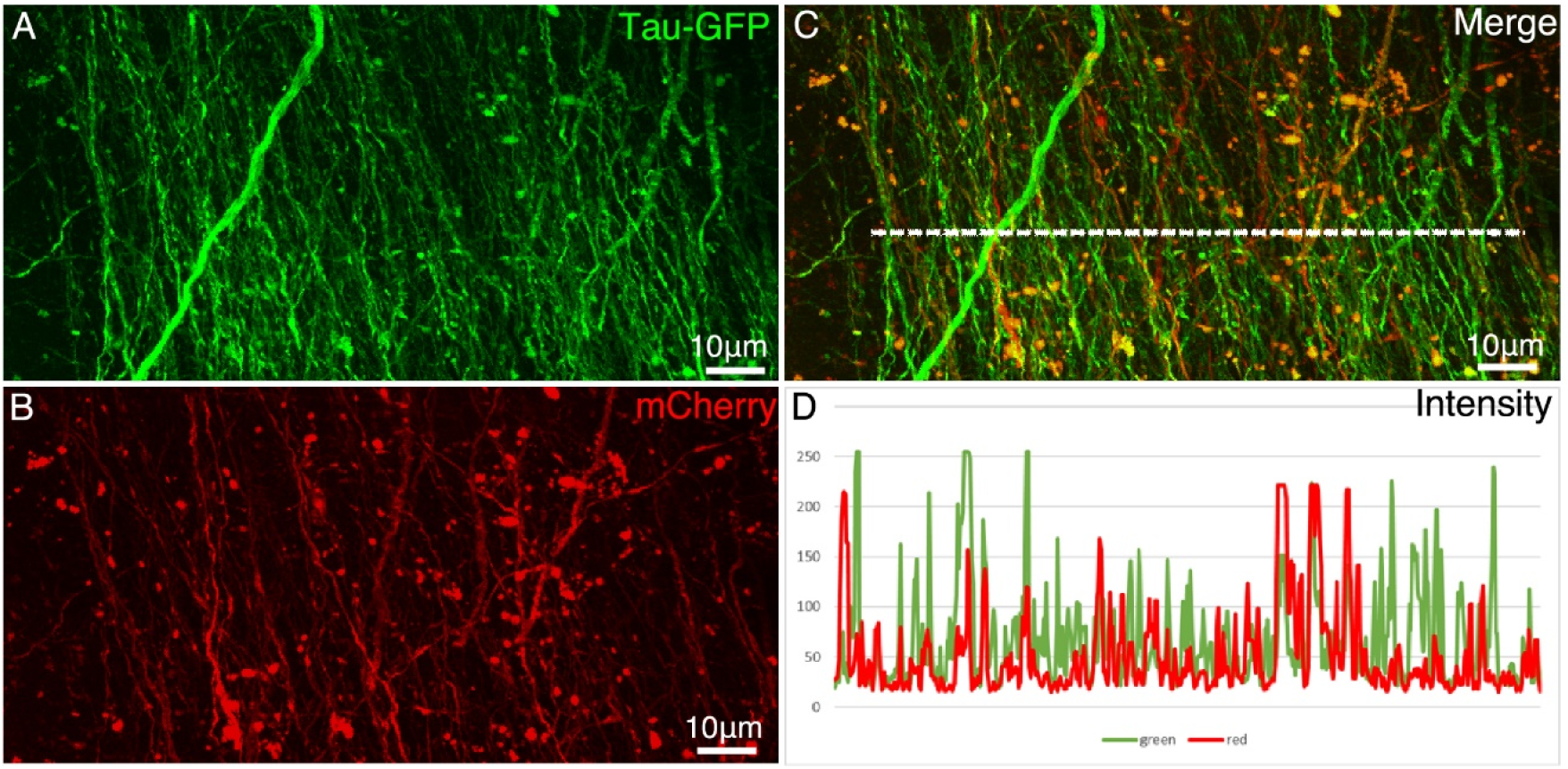
Comparison of two AAV constructs. AAV2/9 encoding Tau-GFP and AAV2/9 encoding mCherry were co-injected in the premotor cortex. Figures **A** and **B** show the axon fibers labeled by Tau-GFP and mCherry, respectively. (**C**) Colocalization of mCherry and GFP in the axonal fibers. (**D**) The intensity profiles (measured using ImageJ on 8-bit TIF images) along the dashed line (in C) in red and green channels. After normalization, a direct comparison indicates that the intensity of Tau-GFP was stronger than that of mCherry.

**Fig. S4.**
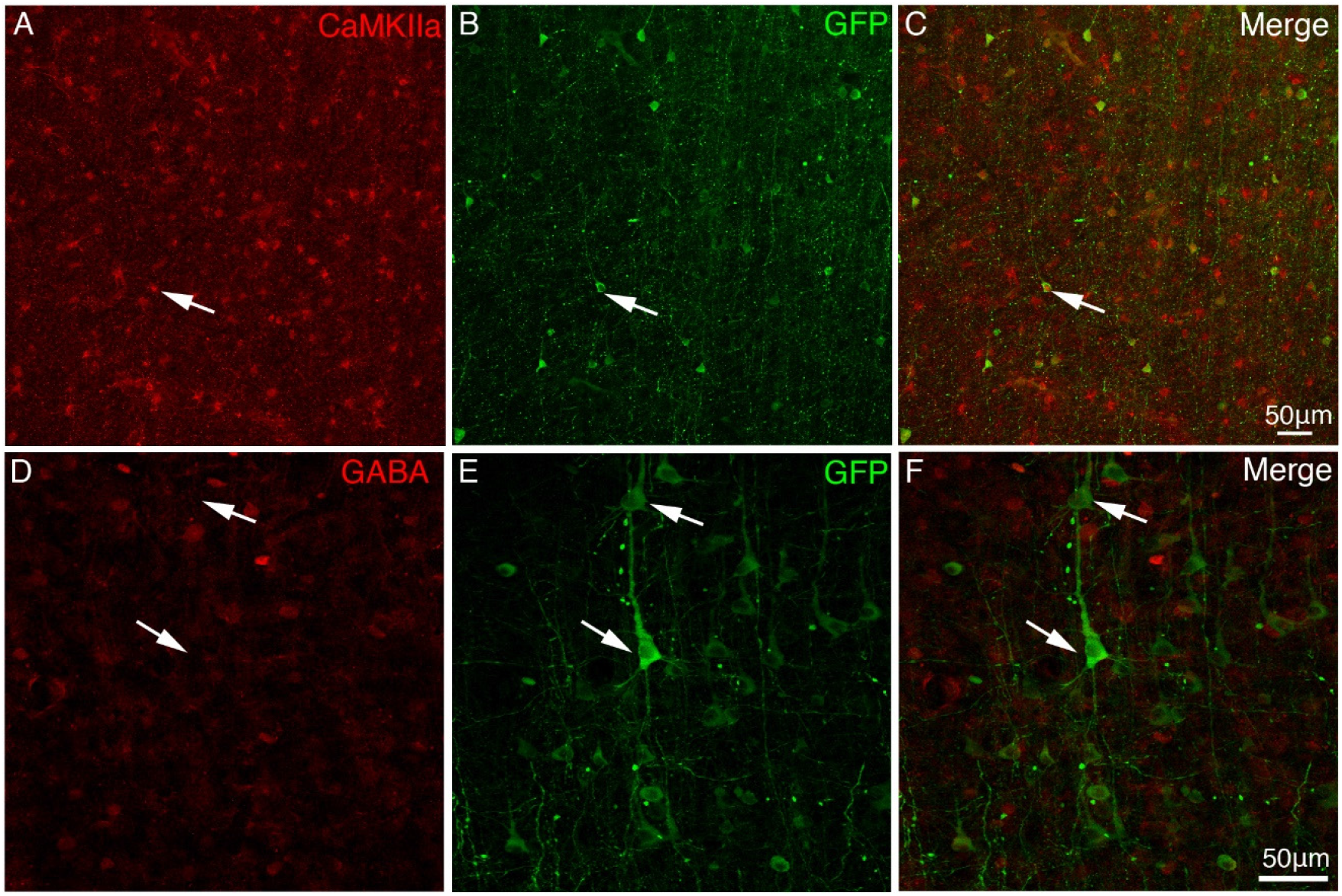
Expression of Tau-GFP in excitatory neurons in macaque brain. To identify neuronal transduction of AAV2/9 constructed with CaMKIIα promoter, cortical sections of vlPFC were immunostained with anti-CaMKIIα (A-C) and anti-GABA (D-F) antibodies. (**A**-**C**) Tau-GFP-expressing neurons around the injection site were identified CaMKIIα positive. (**D**-**F**) Immunofluorescent staining with GABA antibody shows that the GFP-expressing neurons were negative with GABA. Arrowheads indicate Tau-GFP positive cell bodies.

**Fig. S5.**
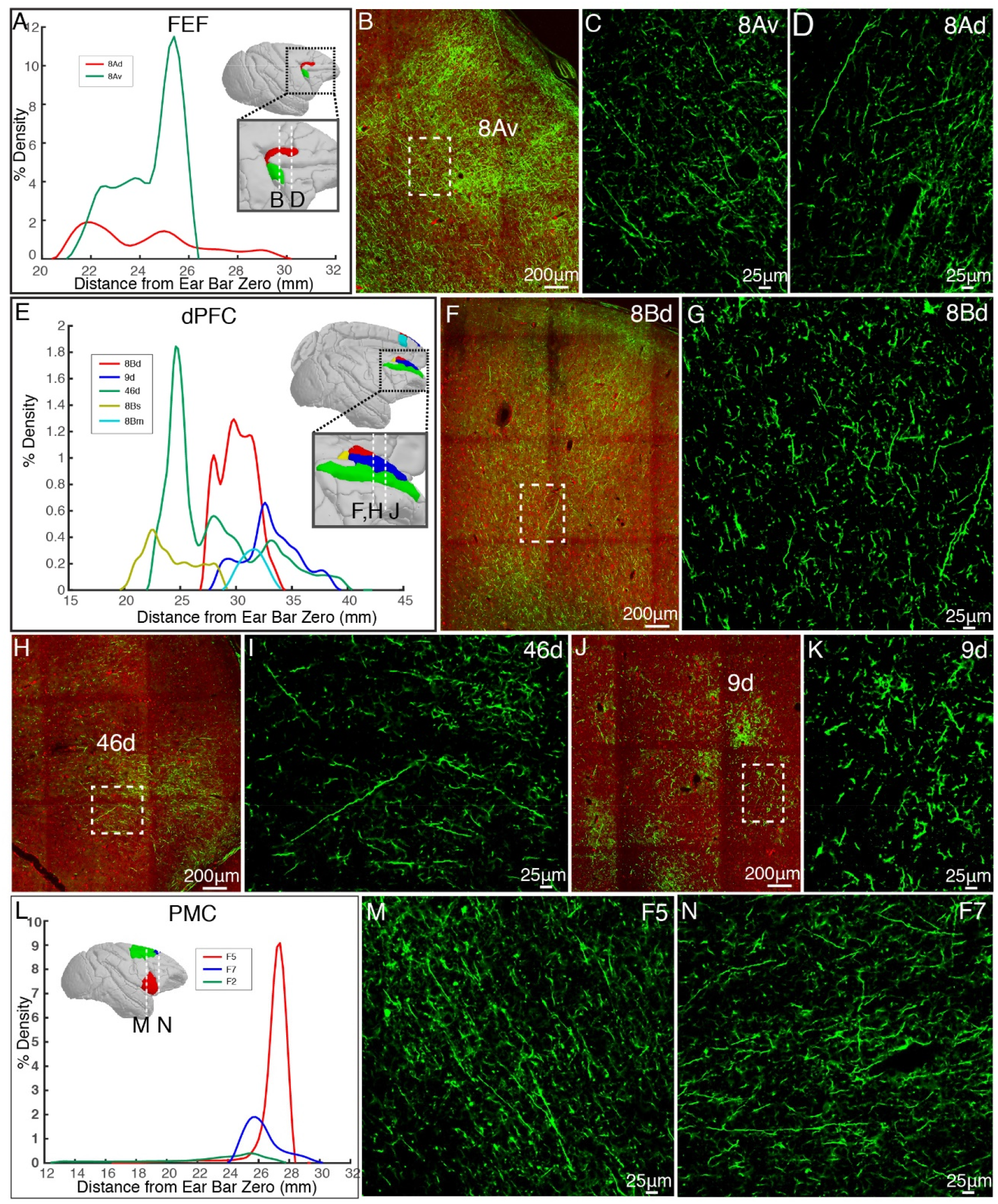
Representative cortical projections of vlPFC. (**A**) Percentage of output density of vlPFC projectome along the anterior-posterior axis of the FEF. (**B**-**D**) Representative two-photon images of vlPFC axonal projections to FEF: 8Av and 8Ad. (**E**) Percentage of output density of vlPFC projectome along the anterior-posterior axis of the dPFC. (**F**-**J**) Representative two-photon images of vlPFC axonal projections to dPFC: 8Bd, 46d, and 9d. (**L**) Percentage of output density of vlPFC projectome along the anterior-posterior axis of the PMC. (**M**, **N**) Representative images of vlPFC axonal projections to PMC: F5 and F7.

**Fig. S6.**
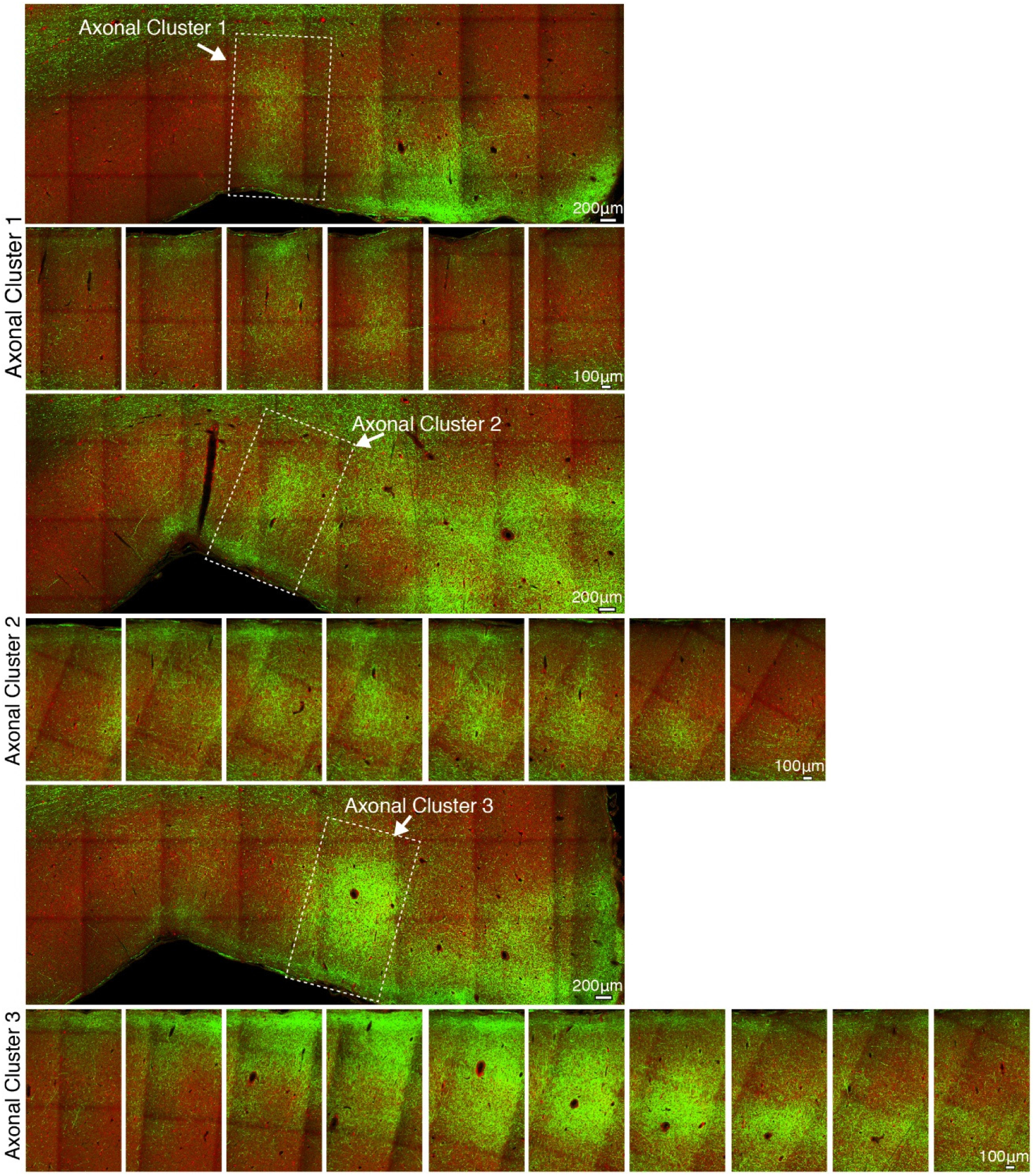
The spatial extent of axonal clusters in the frontal cortex. GFP-labeled axonal fibers issued from vlPFC are overlaid with the autofluorescence image (acquired through the red channel) which provides anatomical landmarks of brain structures. Three representative axonal clusters are shown along the z-axis extent at 200 μm interval between each slice.

**Fig. S7.**
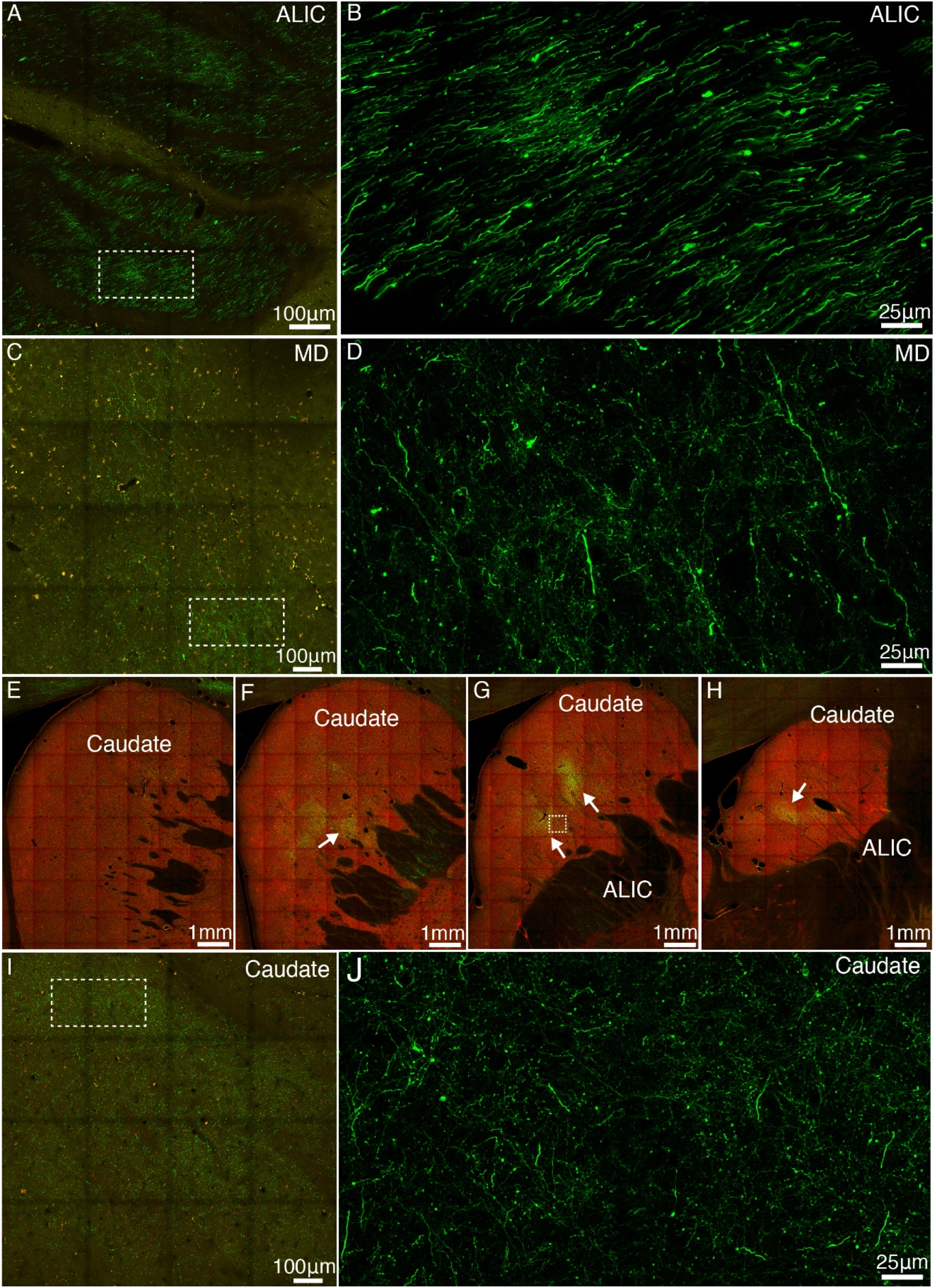
Representative subcortical projections of vlPFC. (**A**) GFP-labeled axonal bundles pass through the anterior limb of internal capsule (ALIC). The image of green channel was superimposed on to that of red channel for better visualization. (**B**) Close-up view of the boxed region in A. (**C**) GFP-labeled axonal fibers in the mediodorsal (MD) thalamus. (**D**) Close-up view of the boxed region in C. (**E**-**H**) GFP-labeled axon clusters (indicated by white arrows) were found in the medial and caudal parts of caudate. (**I**-**J**) Close-up view of the dashed box in g with two levels of magnification.

**Fig. S8.**
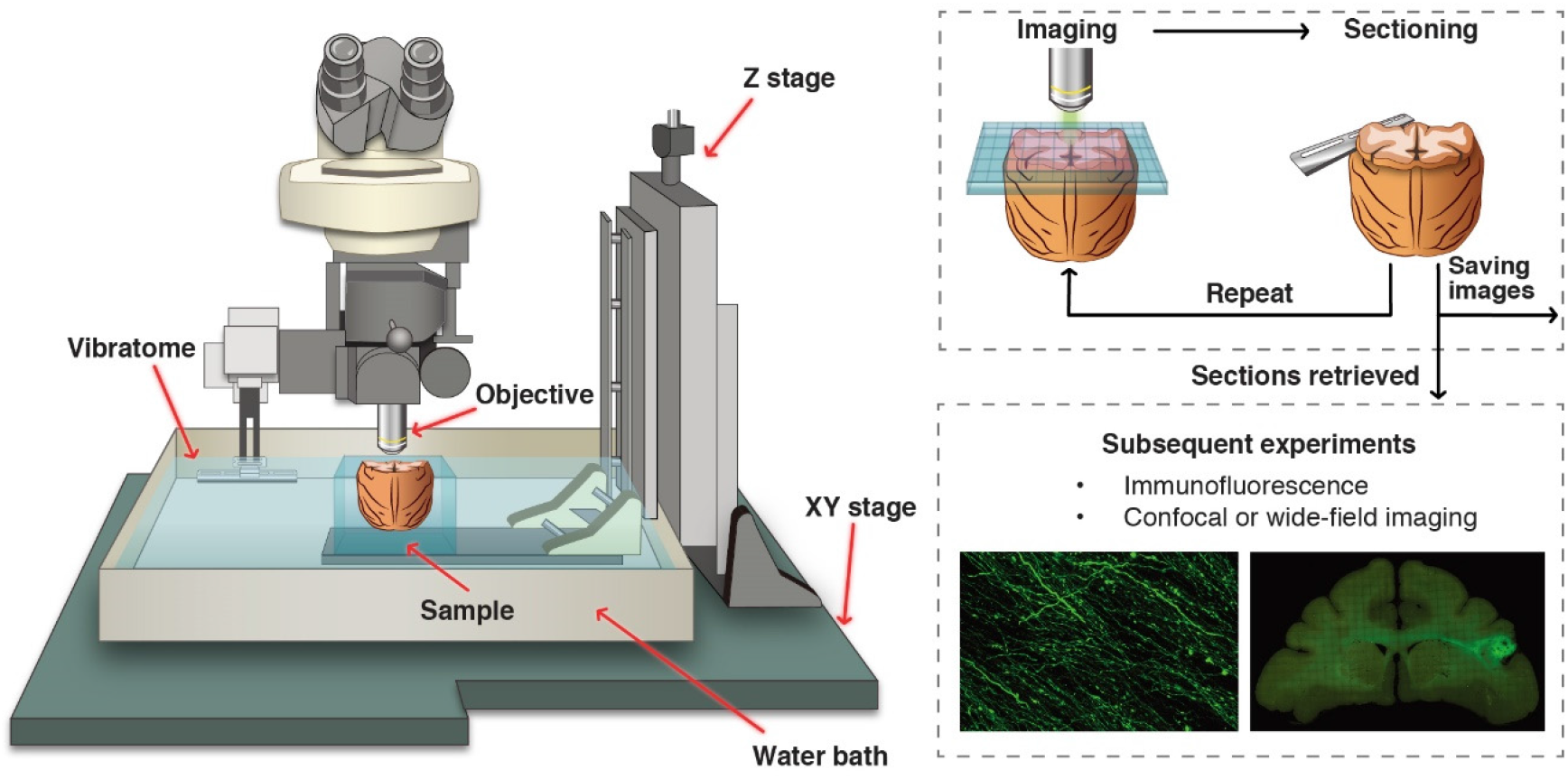
A customized setup of serial two-photon tomography for macaque brain. The STP tomography was integrated with a two-photon microscope and a vibratome, both of which were computer controlled and fully automated. The traveling range of XYZ stage was customized to 50 by 60 by 70 mm to cover the entire macaque brain.

**Fig. S9.**
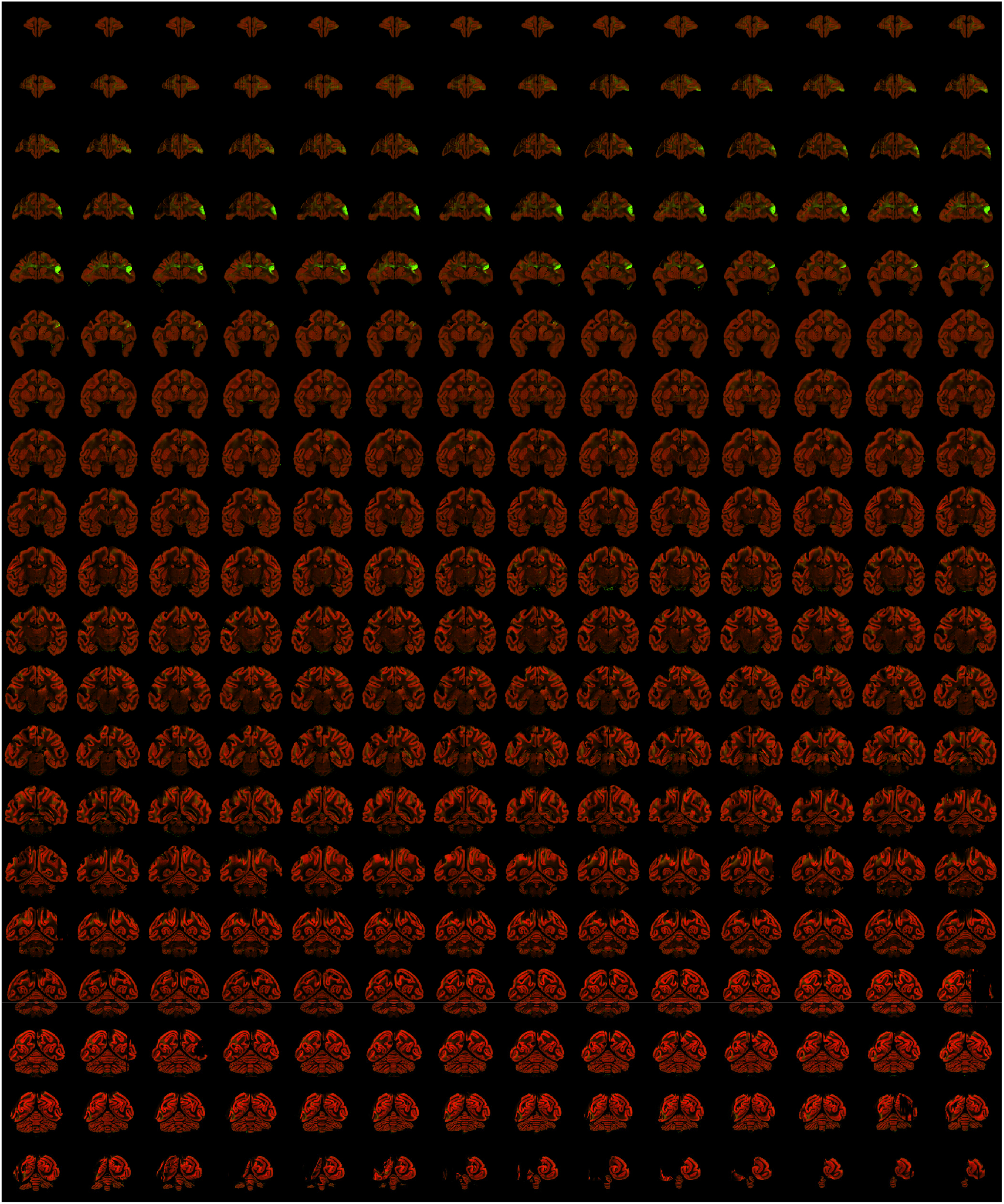
A typical STP tomography image set of a single macaque brain. High-resolution (0.95 μm/pixel) images of serial sections of a single brain are shown as an example (injection into vlPFC). The stepping distance along z-axis is 200 μm. The images acquired through the red channel are used as background. The injection site (vlPFC) and major projection targets can be readily observed in this ‘montage’ view.

**Fig. S10.**
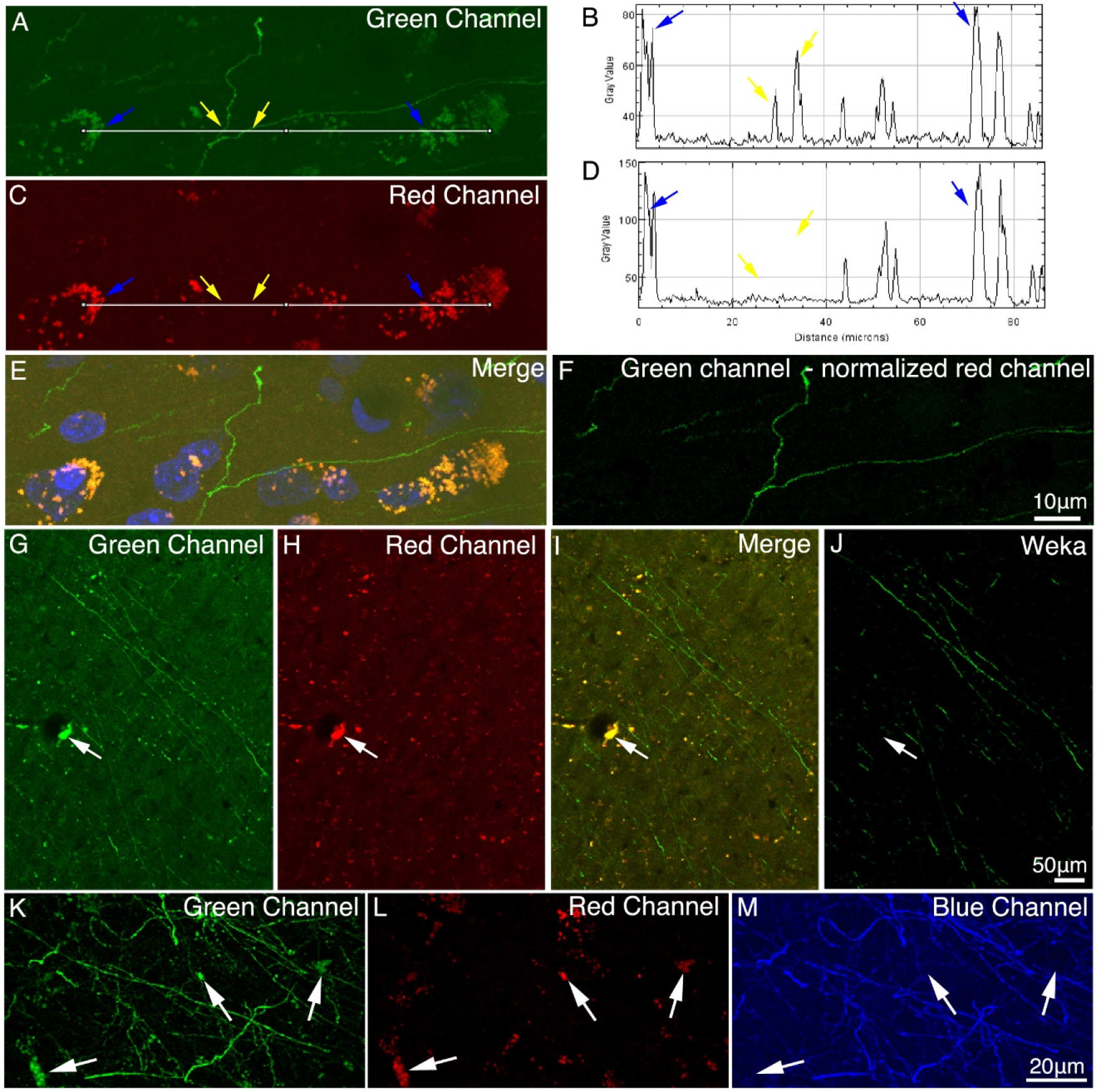
Methods for background reduction. (**A**-**D**) Properties of autofluorescence in macaque brain. Images in the right panel (B and D) show intensity profiles (measured using ImageJ on 8-bit TIF images) along the white line in the left panel (A and C). Yellow arrows indicate GFP signal, and blue arrows indicate background autofluorescence. Due to the broad emission spectrum of autofluorescence (**E**), the autofluorescence was reduced by subtracting the normalized autofluorescence signal in the red channel from the green channel (**F**). (**G**-**J**) A supervised machine learning approach was applied for removing autofluorescence. The classifier was trained to distinguish the GFP signal from background, and the segmented image (J) contained only GFP signal. White arrows indicate the autofluorescence puncta in image G-I. (**K**-**M**) The brain slice was stained with anti-GFP antibody and Alexa Fluor 405 conjugated secondary antibody to convert the GFP signal from green channel to blue channel. Unlike the green (K) and red (L) channels, the converted blue channel (M) did not contain high intensity autofluorescence puncta.

**Fig. S11.**
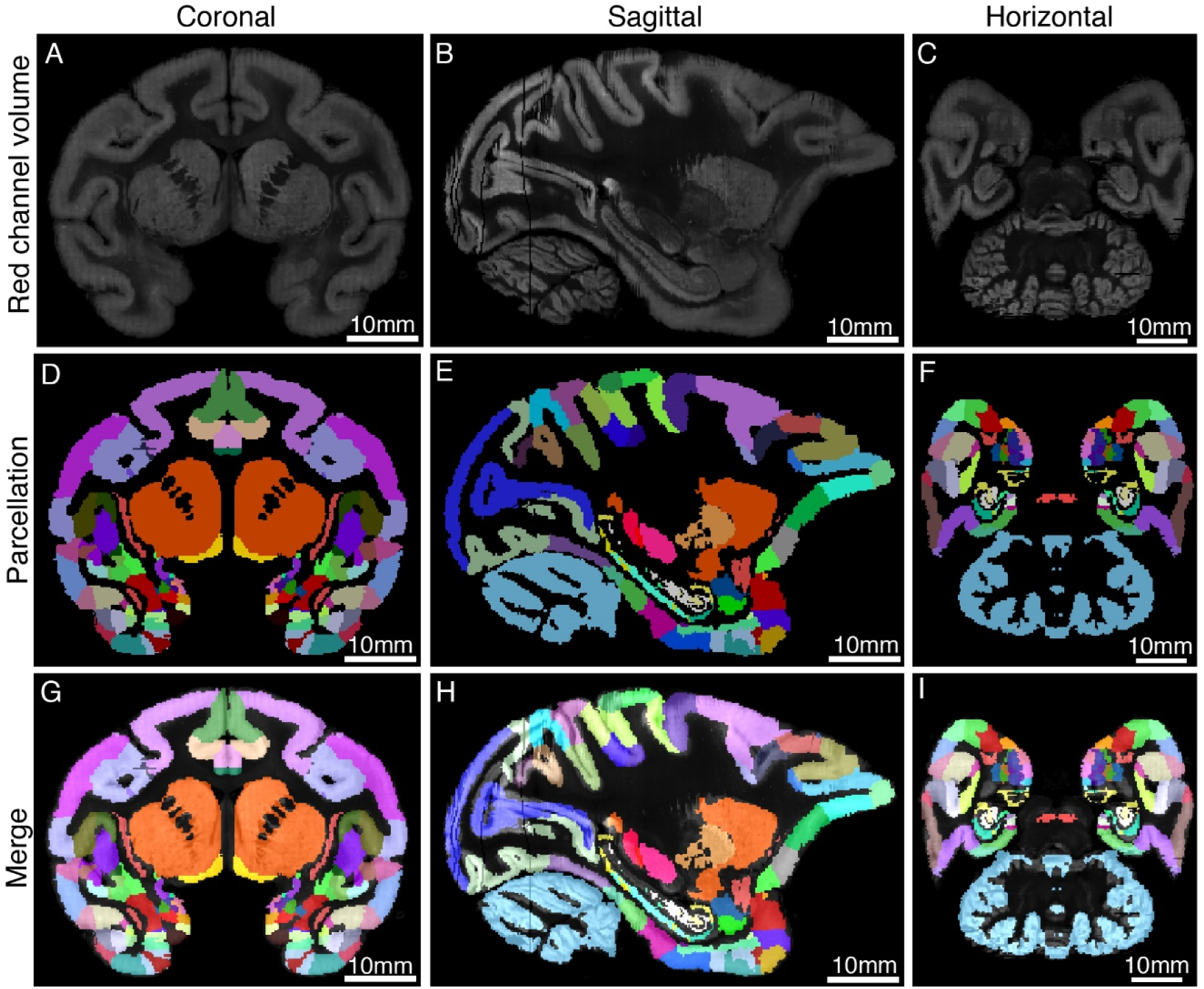
Co-registration of STP images to the MRI-based template of macaque brain. The STP tomography volume and the T1 volume of the monkey brain template were co-registered in a common 3D space. The upper panel shows the coronal (**A**), sagittal (**B**), and horizontal (**C**) images of the warped red channel of STP tomography. (**D**-**F**) The parcellation of Saleem and Logothetis atlas was displayed on three planes. (**G**-**I**) The fluorescence images of STP data were merged with the MRI atlas of macaque brain.

